# GLP-1 Notch - LAG-1 CSL control of the germline stem cell fate is mediated by transcriptional targets *lst-1* and *sygl-1*

**DOI:** 10.1101/825232

**Authors:** Jian Chen, Ariz Mohammad, Nanette Pazdernik, Huiyan Huang, Beth Bowman, Eric Tycksen, Tim Schedl

**Affiliations:** Department of Genetics, Washington University School of Medicine; Department of Pediatrics, Washington University School of Medicine; Department of Biology, Emory University; Genome Technology Access Center, McDonnell Genome Institute, Washington University School of Medicine; Integrated DNA Technologies; Vanderbilt University

## Abstract

Stem cell systems are essential for the development and maintenance of polarized tissues. Intercellular signaling pathways control stem cell systems, where niche cells signal stem cells to maintain the stem cell fate/self renewal and inhibit differentiation. In the *C. elegans* germline, GLP-1 Notch signaling specifies the stem cell fate. We undertook a comprehensive genome-wide approach to identify transcriptional targets of GLP-1 signaling. We expected primary response target genes to be evident at the intersection of genes identified as directly bound by LAG-1, the *C. elegans* Notch pathway sequence-specific DNA binding protein, from ChIP-seq experiments, with genes identified as requiring GLP-1 signaling for RNA accumulation, from RNA-seq analysis. Furthermore, we performed a time-course transcriptomics analysis following auxin inducible degradation of LAG-1 to distinguish between genes whose RNA level was a primary or secondary response of GLP-1 signaling. Surprisingly, only *lst-1* and *sygl-1*, the two known target genes of GLP-1 in the germline, fulfilled these criteria, indicating that these two genes are the primary response targets of GLP-1 Notch and may be the sole germline GLP-1 signaling protein-coding transcriptional targets for mediating the stem cell fate. In addition, three secondary response genes were identified based on their timing following loss of LAG-1, their lack of a LAG-1 ChIP-seq peak and that their *glp-1* dependent mRNA accumulation could be explained by a requirement for *lst-1* and *sygl-1* activity. Moreover, our analysis also suggests that the function of the primary response genes *lst-1* and *sygl-1* can account for the *glp-1* dependent peak protein accumulation of FBF-2, which promotes the stem cell fate and, in part, for the spatial restriction of elevated LAG-1 accumulation to the stem cell region.

**Author Summary:** Stem cell systems are central to tissue development, homeostasis and regeneration, where niche to stem cell signaling pathways promote the stem cell fate/self-renewal and inhibit differentiation. The evolutionarily conserved GLP-1 Notch signaling pathway in the *C. elegans* germline is an experimentally tractable system, allowing dissection of control of the stem cell fate and inhibition of meiotic development. However, as in many systems, the primary molecular targets of the signaling pathway in stem cells is incompletely known, as are secondary molecular targets, and this knowledge is essential for a deep understanding of stem cell systems. Here we focus on the identification of the primary transcriptional targets of the GLP-1 signaling pathway that promotes the stem cell fate, employing unbiased multilevel genomic approaches. We identify only *lst-1* and *sygl-1*, two of a number of previously reported targets, as likely the sole primary mRNA transcriptional targets of GLP-1 signaling that promote the germline stem cell fate. We also identify secondary GLP-1 signaling RNA and protein targets, whose expression shows dependence on *lst-1* and *sygl-1*, where the protein targets reinforce the importance of posttranscriptional regulation in control of the stem cell fate.

## Introduction

Stem cell systems are required for the development and maintenance of polarized tissues, controlling the position, number and timing of differentiated cell type production. Stem cell systems have non-stem niche cells that signal nearby cells to promote the stem cell fate/self renewal and to inhibit differentiation. Niche – stem cell signaling pathways include Notch, BMP, Wnt and JAK/Stat [1–3]. A deep understanding of how a stem cell system works requires knowledge of the full repertoire of genes that are the primary targets of the signaling pathway, as well as secondary response gene products.

Here we focus on the *C. elegans* germline stem cell system, which shares a number of features with other stem cell systems [4]. Niche - germline stem cell signaling employs the Notch pathway, which has been extensively studied in *C. elegans* [5]. The worm germline is a polarized tube-shaped cellular assembly line designed for the rapid production of gametes. The germline is capped by the distal tip cell (DTC), which is a large somatic niche cell that polarizes germline cellular organization. Germ cells adjacent to the DTC are in a region called the progenitor zone, which distally contains stem cells, then cells completing a terminal mitotic cell cycle and cells undergoing meiotic S-phase, followed proximally by cells undergoing the earliest stages of meiotic prophase, leptotene and zygotene [4,6,7](**Fig 1**). The DTC expresses two Notch pathway DSL (for Delta, Serate, LAG-2) ligands, LAG-2 and APX-1 [8–10]. *C. elegans* has two Notch receptors, GLP-1 and LIN-12 [5, 11]. GLP-1 is expressed in progenitor zone germ cells and continuously required to promote the germline stem cell fate. Genetic loss of *glp-1* in larval or adult stages results in loss of all germline stem cells because of their premature entry into meiotic prophase [12, 13]. Conversely, <u>g</u>ain of function (gf) mutations in *glp-1* result in a tumorous germline, with a vast excess of stem cells and reduced or no meiotic prophase cells [14, 15]. *lin-12* functions only in somatic cell fate specification, including mediating the anchor cell – ventral uterine decision and 2° vulval cell fate specification [16]. *lin-12* and *glp-1* function redundantly in late embryonic fates, and their simultaneous loss results in an early larval lethal phenotype called Lag (for *lin-12* <u>a</u>nd *<u>g</u>lp-1*; [11]). The current model is that when the DTC presenting LAG-2 and APX-1 interacts with GLP-1 expressed in germ cells, ligand dependent cleavage of the receptor generates the GLP-1 intra<u>c</u>ellular <u>d</u>omain, GLP-1(ICD), which translocates to the nucleus and associates with the sequence specific DNA binding protein LAG-1 to activate primary target gene transcription. LAG-1 is a founding member of the CSL family of DNA binding proteins [for <u>C</u>BF1 (or RBPJ) in mammals, <u>S</u>u(H) in Drosophila and <u>L</u>AG-1] [17]. Complete loss of *lag-1* results in L1 larval lethality, reflecting LAG-1 functioning as the DNA binding co-factor for both GLP-1(ICD) and LIN-12(ICD), while a *lag-1* hypomorphic mutant results in incompletely penetrant loss of germline stem cells due to premature entry into meiotic prophase [11, 18]. LAG-1 and other CSL proteins share the same *in vitro* DNA binding site (GTGGGAA, LAG-1/CSL binding site hereafter) [17,19,20]. GLP-1(ICD), bound to LAG-1, and associated with SEL-8 (also called LAG-3) that functions similarly to Drosophila Mastermind, forms an activation complex that transcribes GLP-1 signaling targets. LAG-1 is thus central to GLP-1 signaling as its sequence specific DNA binding determines direct transcriptional targets. Through its transcriptional targets, GLP-1 signaling pathway promotes the stem cell fate, at least in part, by inhibiting three parallel pathways that promote meiotic development, the GLD-1 pathway, the GLD-2 pathway and SCF^PROM-1^ (**Fig 1**).

**Figure 1.**
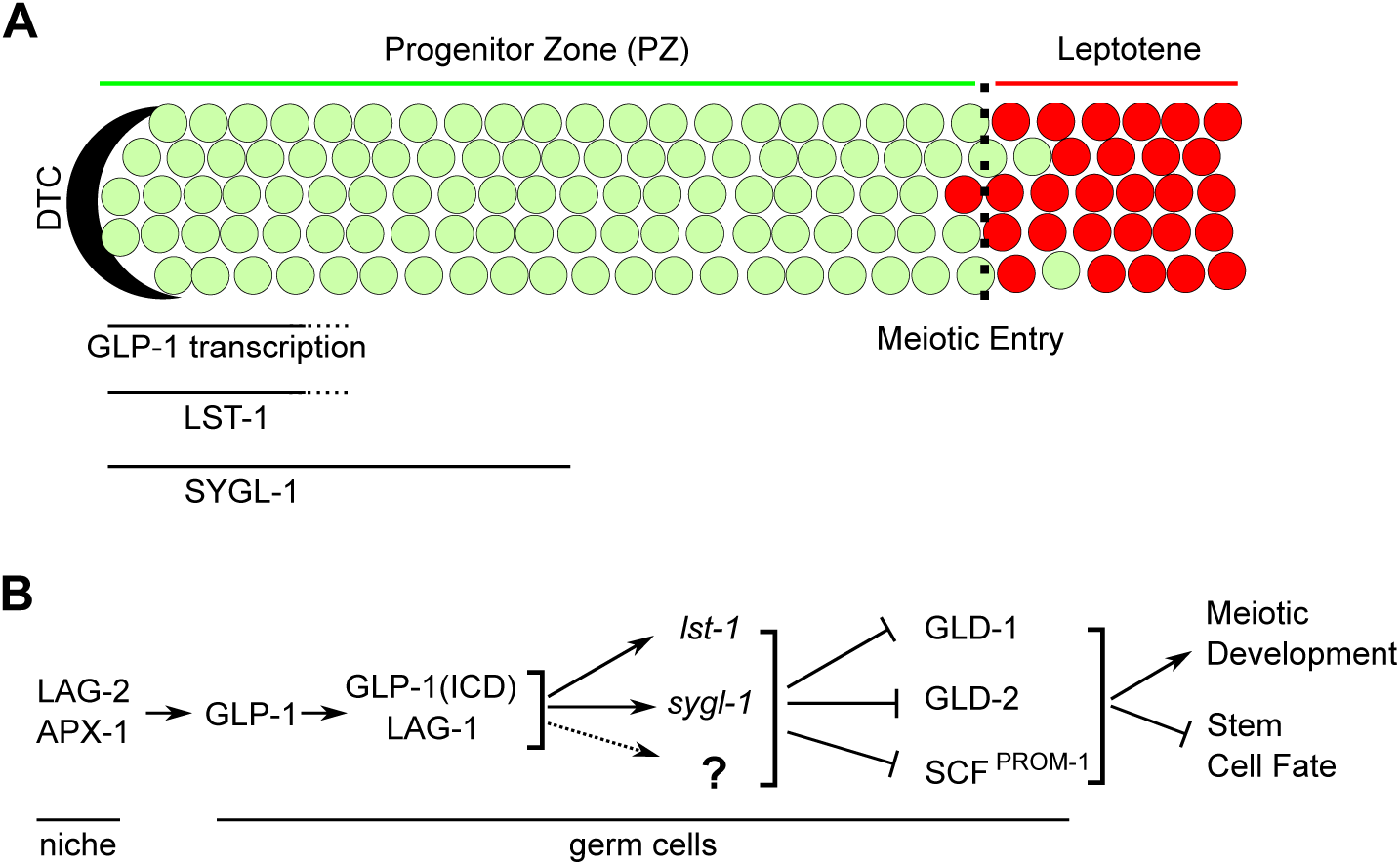
Overview of GLP-1 signaling in the distal germline of *C. elegans*. (A) Schematic of the adult hermaphrodite distal germline. The distal end of the germline is capped by a somatic distal tip cell (DTC). Progenitor zone cells are shown in green; meiotic prophase cells are in red. Dashed line indicates the progenitor zone – meiotic prophase boundary, the operationally defined point of meiotic entry [4]. GLP-1 signaling maintains germline stem cell fate and GLP-1 mediated transcription occurs in the distal most ∼5 cell diameter (cd) of the germline [22, 42]. LST-1 and SYGL-1 protein are observed in the distal most 5 or 10 cd, respectively [23]. (B) Genetic pathway controlling the stem cell fate vs meiotic development decision in the distal end of *C. elegans* germline. GLP-1 signaling acts, at least in part, through transcriptional targets *lst-1* and *sygl-1* to repress the GLD-1, GLD-2 and SCF^PROM-1^ meiotic entry pathways (reviewed in [4]). GLP-1(ICD), GLP-1 intracellular domain.

Two germline GLP-1 signaling transcriptional targets, *lst-1* and *sygl-1*, have been identified that are redundantly required to promote the stem cell fate through a candidate gene approach [21, 22]. A number of lines of evidence support this identification. First, genetic manipulation of *lst-1* and *sygl-1* give the same phenotypes as manipulation of *glp-1*: the *lst-1 sygl-1* double null mutant has the identical premature meiotic entry of all cells in the L2 stage as *glp-1* null; loss of *lst-1* and *sygl-1* later in larval and early adult stages results in all stem cells entering meiotic prophase as is observed with loss of *glp-1*; and ubiquitous overexpression of *lst-1* or *sygl-1* in the germline results in a tumorous phenotype similar to *glp-1(gf)* [21, 23]. Second, epistasis analysis place *lst-1* and *sygl-1*, like *glp-1*, upstream of the meiotic entry pathway genes. Third, *lst-1* and *sygl-1* nascent transcripts are restricted to the distal most germ cells, contacting the DTC, and this expression requires *glp-1* activity [21, 22] (**Fig 1**). Fourth, LAG-1 binding sites in a *sygl-1* promoter reporter are required for distal germline specific expression [21]. Finally, *lst-1* was previously identified as a LIN-12 transcriptional target involved in vulval development [24]. LST-1 and SYGL-1 act, at least in part, through binding to Pumilio family RNA binding proteins FBF-1 and FBF-2 that function in mRNA degradation and translational repression of the *gld-1* meiotic entry pathway gene [23,25–28].

*lst-1* and *sygl-1* were identified in a small candidate RNAi screen of 15 genes that meet the following criteria – (i) the gene contained a cluster of at least four LAG-1 binding sites and (ii) its mRNA was a target of FBF-1 regulation based on FBF-1 immunoprecipitation followed by microarray analysis [21,24,29]. The presence of LAG-1 binding sites in genes regulated by Notch signaling transcription is logically predicted. However, as FBF-1 promotes the stem cell fate and functions in mRNA degradation and translational repression, it is counterintuitive that GLP-1 signaling transcriptional targets would be FBF-1 posttranscriptional repression targets. In addition, three other GLP-1 signaling germline transcriptional targets have been reported, *fbf-2*, *utx-1* and *lip-1* [30–32]. Despite these findings, the full repertoire of the GLP-1 signaling primary transcriptional targets is unknown.

We have taken an unbiased genome-wide approach to identify primary transcriptional targets of GLP-1 signaling, through intersection of genes identified as directly bound by LAG-1, from ChIP-seq experiments, with genes identified as requiring GLP-1 signaling for RNA accumulation, from RNA-seq transcriptomics analysis. An important part of our approach was identifying genes whose RNA level, or protein level, was dependent on *glp-1* signaling. However, in *glp-1* null mutants, germ cells enter meiosis in the L2 stage complicating comparative RNA and protein accumulation studies because of very different germ cell number and type. Therefore, we have taken advantage of the epistasis of *gld-2 gld-1* double null over the *glp-1* null mutant, comparing meiotic entry defective tumorous germlines, with and without *glp-1* activity, similar to other studies [21–23,30,32,33]. We identified *lst-1* and *sygl-1* as the only genes that had both germline LAG-1 binding and whose expression was dependent on *glp-1* and germline *lag-1* activity. Furthermore, we used a time course transcriptomic analysis following auxin-induced degradation of LAG-1 to distinguish between primary versus secondary changes in RNA level in GLP-1 signaling. *lst-1* and *sygl-1* were the first genes (2hrs treatment) whose mRNA level were dependent on LAG-1 activity, representing a primary effect. Five additional genes were then identified at a later time point (4hrs treatment) whose RNA accumulation was dependent of LAG-1 activity. Consistent with these changes being a secondary effect, the five genes were not bound by LAG-1 and for three of the genes, RNA accumulation was dependent on *lst-1* and *sygl-1* activity. We additionally showed that *glp-1* dependent peak FBF-2 accumulation [30] is fully explained by a requirement for downstream *lst-1* and *sygl-1* activity. Finally, we expand our understanding of *lag-1* function and expression. We showed that *lag-1* is germline autonomously required for the stem cell fate and found that LAG-1 is spatially restricted, with peak accumulation in the stem cell region of the progenitor zone. Peak accumulation is regulated posttranscriptionally, with about 50% of *glp-1* dependent LAG-1 accumulation contributed by *lst-1* and *sygl-1* activity. Together, our results are consistent with the possibility that *lst-1* and *sygl-1* are the only mRNA transcriptional targets and that GLP-1 signaling is mediated largely or completely by *lst-1* and *sygl-1*.

## Results

### LAG-1 expression is spatially restricted to stem cells and autonomously required in the germline for the stem cell fate

As a first step in examining GLP-1 Notch – LAG-1 CSL transcriptional control of the germline stem cell fate, we determined the LAG-1 protein accumulation pattern. The endogenous *lag-1* locus was tagged with 3xHA at the C-terminus of the LAG-1, using CRISPR/Cas9 engineering, to generate *lag-1(oz530[lag-1::3xHA])*, hereafter called *lag-1::HA* (**Fig 2A; Materials and Methods**). The *lag-1:: HA* strain appears phenotypically wild type (**S1 Fig**). We examined germline LAG-1::HA accumulation by anti-HA antibody staining in dissected hermaphrodite gonad preparations (**Fig 2B-C; Materials and Methods**). In young adults (1 day past mid-L4 larval stage) LAG-1::HA was found in germ cell nuclei in the distal most ∼10 <u>c</u>ell <u>d</u>iameters (cd hereafter) of the progenitor zone (PZ), during late pachytene, diplotene and diakinesis of oogenesis. The late oogenic accumulation is consistent with maternal loading of LAG-1 for early embryonic GLP-1 signaling in specification of certain blastomere cell fates [34, 35]. In mid/late L4, LAG-1::HA was also found in germ cell nuclei in the distal most ∼10 cd of the PZ, but not observed in late pachytene cells in the proximal gonad, which correspond to germ cells undergoing spermatogenesis [36]. Distal germline LAG-1::HA staining was variable in 1-day adults, and the staining intensity was weaker than in the L4 stage. We also observed strong LAG-1::HA staining in nuclei of all somatic gonad cells, the DTC, and all sheath and spermathecal cells, as well as in polyploidy intestinal cells, in both the L4 and adult stages dissection preparations.

**Figure 2.**
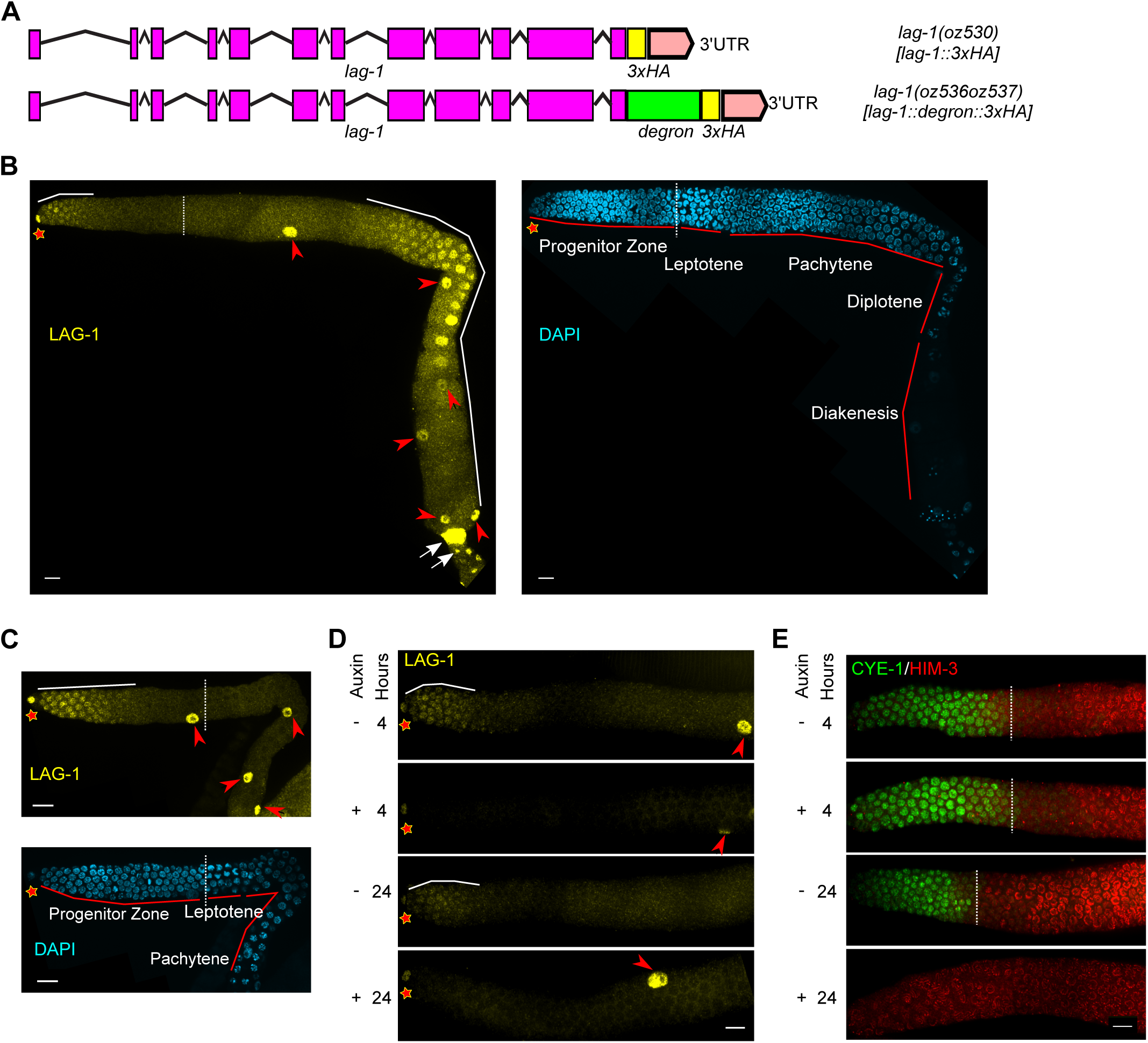
LAG-1 spatial accumulation and germline autonomous function to promote the stem cell fate. (A) Diagrams of tagged endogenous *lag-1* alleles, *lag-1(oz530[lag-1::3xHA])* (top) and *lag-1(oz536oz537[lag-1::degron::3xHA])* (bottom). Purple boxes, exons; lines, introns; pink boxes, untranslated region; yellow boxes, 3xHA; green box, degron. (B & C) Images of HA-stained (LAG-1, yellow) germlines from dissected hermaphrodite gonads, co-stained with DAPI (cyan) for (B) 1-day adult and (C) mid-L4 stage. (D & E) Images of (D) HA-stained (LAG-1, yellow) and (E) CYE-1 (green), HIM-3-stained (red) germlines from dissected hermaphrodites. L4 stage animals with the following genotype *lag-1(oz536oz537[lag-1::degron::3xHA]); ieSi64[gld-1p::TIR1::mRuby::gld-1 3’UTR]* were treated with or without auxin for 4 hours (top two panels) or 24 hours (bottom two panels). Note that depending on the orientation of the gonad when mounted for microscopy, two, one or zero distal sheath cell nuclei are visible in the surface views of the germline shown in the photographs. Asterisk, distal end; dashed lines, position of meiotic entry; solid white lines, position of LAG-1 accumulation; red arrowhead, sheath cell nuclei; white arrows, spermatheca nuclei. Scale bar is 10 μm.

Knowledge of the cell type in which LAG-1 promotes the germline stem cell fate is a prerequisite for genome-wide studies. While LAG-1 has been assumed to function in the germline to promote the stem cell fate, observing strong LAG-1 accumulation in the DTC raises the possibility that it may have a non-autonomous function in promoting the stem cell fate. We used the auxin-inducible degradation (AID) system that destabilizes degron tagged proteins [37], to test if LAG-1 is require in the germline to promote the stem cell fate. The degron tag, followed by 3xHA, was placed at the C-terminus of LAG-1 in the endogenous locus using CRISPR/Cas9 engineering, to generate *lag-1(oz536oz537)*, hereafter called *lag-1::degron::HA* (**Fig 2A; Materials and Methods**). Germline restricted degradation was achieved by germline specific expression of the TIR1 F-box substrate specificity protein, using the *gld-1* promoter [37] (*ieSi64[gld-1p::TIR1::mRuby]*; **Materials and Methods**). In the absence of auxin treatment, the *lag-1::degron::HA* strain is phenotypically wild type, with LAG-1::degron::HA accumulation indistinguishable from LAG-1::HA (**Fig 2B, D; S1 Fig**). Auxin mediated degradation was initiated at mid-L4. After 4 hrs of auxin treatment, LAG-1::degron::HA distal germ cell nuclear staining was no longer detected and this was also true at 24hrs of auxin treatment (**Fig 2D**). In contrast, strong LAG-1::degron::HA staining remained in the DTC, sheath and spermathecal cells and the intestine at both 4 and 24 hrs of auxin treatment, consistent with germline specific degradation of LAG-1 (**Fig 2D**).

To assess inappropriate entry of stem cells into meiotic prophase following loss of LAG-1::degron::HA, we stained dissected gonads for progenitor zone marker CYE-1 cyclin E [38] and meiotic chromosome axis protein HIM-3 [39, 40] (**Materials and Methods**). After 4 hrs of auxin treatment, when LAG-1::degron::HA is no longer detected, the position of the progenitor zone – leptotene boundary [in cd from the distal tip] was not significantly different from wild type and from animals not treated with auxin (**Fig 2E; S1 Fig**). Following 24 hrs of auxin treatment all progenitor zone cells were CYE-1 negative and HIM-3 positive, indicating that all the stem cells had entered meiotic prophase. The kinetics of stem cell meiotic entry following germline loss of LAG-1::degron::HA (also see below) are consistent with the kinetics observed following loss of GLP-1 signaling through shift of *glp-1* temperature sensitive mutants to the restrictive temperature; the progenitor zone is maintained at 4 hrs and absent by 10/12 hrs [41]. Together, the above results indicate that LAG-1 is germline autonomously required and expressed in the appropriate distal germ cells to promote the stem cell fate.

### LAG-1 accumulation is positively regulated by GLP-1 signaling at a posttranscriptional level and negatively regulated by the GLD-1 and GLD-2 meiotic entry pathways

We next investigated how LAG-1 accumulation is regulated in the distal germline. The *lag-1* gene contains multiple consensus LAG-1/CSL binding sites [17], and along with the distal germline restricted accumulation pattern, suggests the possibility that LAG-1 accumulation occurs through a positive transcriptional loop via GLP-1 signaling. If the GLP-1(ICD) - LAG-1 complex is regulating *lag-1* transcription, we predicted that (i) *lag-1* mRNA would be spatially restricted to the first ∼5 - 10 cd from the distal tip in wild type, where GLP-1 dependent transcription and cytoplasmic mRNA of known targets *lst-1* and *sygl-1* are observed, and (ii) *lag-1* mRNA accumulation would depend on GLP-1 activity [22, 42]. We performed <u>s</u>ingle <u>m</u>olecule fluorescent *in <u>s</u>itu* <u>h</u>ybridization (smFISH) on dissected gonads with *lag-1* mRNA complementary probes, analyzing expression quantitatively by counting foci along the distal-proximal axis of the germline [43](**Materials and Methods**). In wild type, we found that *lag-1* mRNA was uniformly distributed throughout the distal 25 cd; abundant *lag-1* mRNA was found from 10 - 25 cd from the distal tip, a region where *lst-1* and *sygl-1* mRNAs are not observed (**S2 Fig**). To examine GLP-1 dependence, we used the *gld-2 gld-1* null double mutant meiotic entry defective background (**Introduction**) and compared foci distribution relative to wild type in *gld-2 gld-1* with either *glp-1(+)* or the *glp-1* null allele *q175*. As an internal control, wild type gonads were co-dissected and smFISH performed together with *gld-2 gld-1* or *gld-2 gld-1; glp-1(q175)*. The *gld-2 gld-1* mutant background had no significant effect on the uniform distribution of *lag-1* mRNA foci relative to wild type. Furthermore, loss of *glp-1* activity had no significant effect on the uniform distribution of *lag-1* mRNA foci in *gld-2 gld-1* (**S2 Fig**). The uniform distal *lag-1* mRNA accumulation, independent of *glp-1* activity, suggests that GLP-1 signaling is not involved in regulation of *lag-1* transcription in the distal germline.

The above results indicate that distal restricted LAG-1 accumulation occurs through a post-transcriptional mechanism. To investigate this mechanism, we first quantified the distal – proximal accumulation of LAG-1::HA from anti-HA antibody staining in an otherwise wild type background, and subtracted background signal from staining of *lag-1(+)* germlines, lacking 3xHA (**Materials & Methods**). Because germline staining for LAG-1::HA in 24 hrs adults was variable, we employed mid/late L4 hermaphrodites where staining is more consistent from gonad to gonad. High LAG-1::HA was observed from 1 – 5 cd from the distal tip followed by a significant fall to a basal level from ∼17 - 25 cd (**Fig 3A-B**). To allow comparisons, the mean peak intensity at 4 cd was set to 100. From peak to proximal base at 25 cd, there was an ∼7-fold drop in LAG-1::HA levels, with accumulation at base significantly above background, ∼15% of peak.

**Figure 3.**
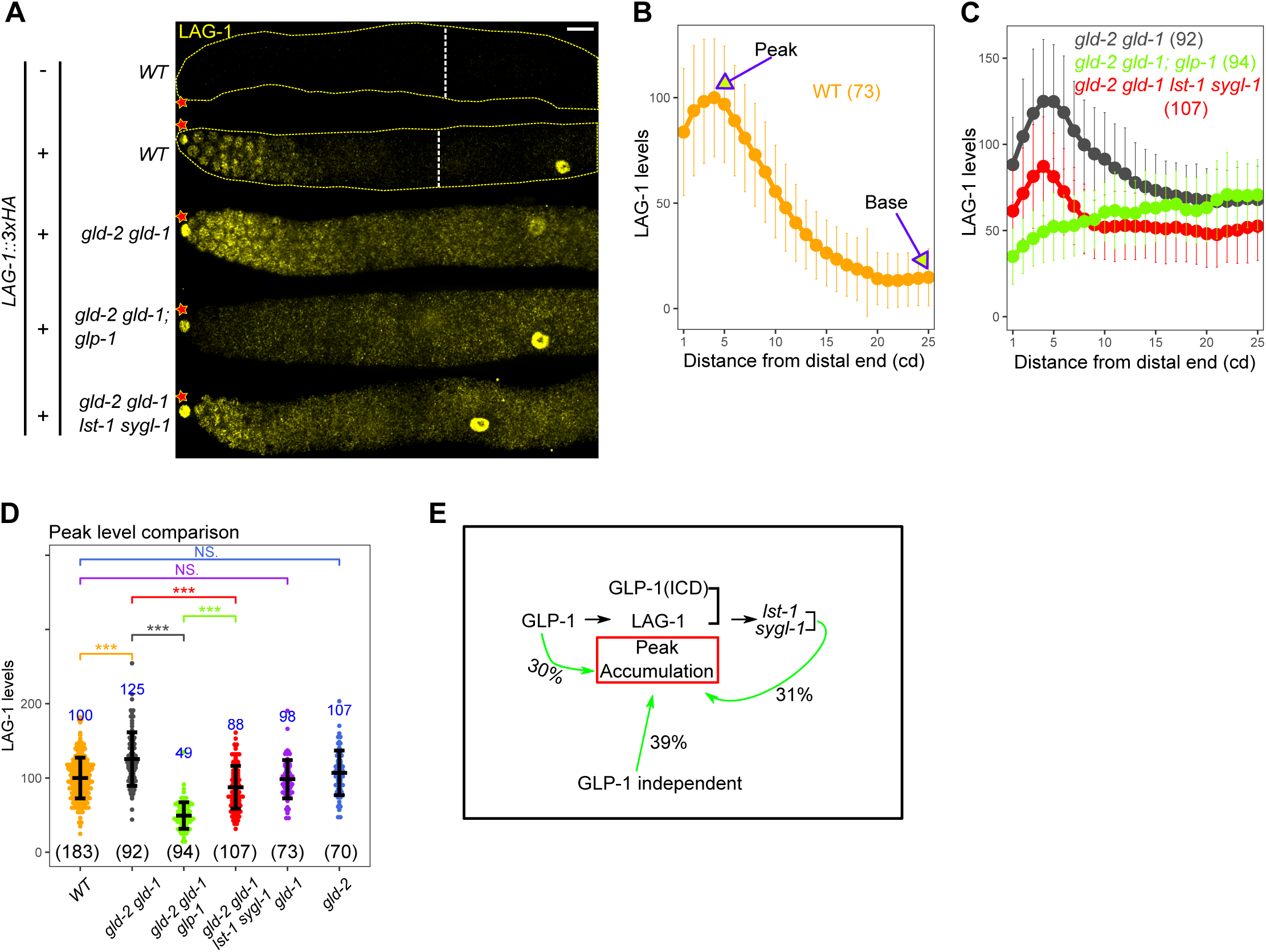
Post-transcriptional regulation of LAG-1 by GLP-1 signaling. (A) Images of HA-stained (LAG-1, yellow) germlines from dissected L4 hermaphrodites of the indicated genotype. See **S1 Table** for the complete genotypes. Asterisk, distal end; dashed white lines, meiotic entry. Scale bar is 10 μm. (B - D) Plot of LAG-1 levels (B - C) and comparison of LAG-1 peak accumulation (D) for indicated genotype. *lag-1(oz530[lag-1::3xHA])* is used for quantification. Numbers indicate mean values of LAG-1 level for each genotype (in blue) and numbers in bracket shows the sample size. Dots, mean (B-C) or data points (D); Error bars, mean ± SD. P-value ≤ 0.01 (*); ≤ 0.001 (**); ≤ 0.0001 (***); > 0.01 non-significant (NS.). (E) Model depicting genetic control of peak accumulation for LAG-1 and percent contribution to peak LAG-1 accumulation. Sixty one percent of LAG-1 peak level is attributed to GLP-1 signaling, in which GLP-1 transcriptional targets *lst-1* and *sygl-1* account for 31% of LAG-1 peak level.

We then investigated the possibility that GLP-1 signaling indirectly regulates LAG-1 peak levels, possibly through targets *lst-1* and *sygl-1*. The pattern and level of LAG-1::HA was first assessed in the *gld-2 gld-1* background, and then in *gld-2 gld-1; glp-1* null triple mutants and the *gld-2 gld-1 lst-1 sygl-1* quadruple null mutants, each co-dissected and stained with wild type, with and without *lag-1::HA* to allow background subtraction and normalization (**Fig 3A, C-D; Materials and Methods**). In *gld-2 gld-1*, LAG-1::HA displayed a similar overall expression pattern as in wild type, but with elevated accumulation throughout the progenitor zone, with peak levels increased by ∼25% (also see below). In the absence of *glp-1*, LAG-1::HA level was reduced throughout the progenitor zone, with a gradual rise from distal to proximal. Compared to peak level at 4 cd, LAG-1::HA is 39% of *glp-1(+)* level, indicating that ∼39% of peak LAG-1 accumulation is independent of GLP-1 signaling, with a similar level continuing into the proximal progenitor zone. Thus, ∼61% of peak LAG-1 accumulation depends on GLP-1 signaling. In the absence of *lst-1* and *sygl-1*, we observed a peak of LAG-1 accumulation that was intermediate between the presence and absence of *glp-1*, ∼70% of the LAG-1::HA peak in *gld-2 gld-1*. The difference in accumulation indicates that *lst-1* and *sygl-1* activity account for ∼50% of *glp-1* dependent LAG-1 accumulation (**Fig 3C-D**). These results do not provide an explanation for the remaining 50% of *glp-1* dependent control of LAG-1 accumulation, which is not reliant on control of *lag-1* mRNA level (**S2 Fig**). Figure 3E summarizes the GLP-1 signaling dependent and independent control of LAG-1 peak accumulation, indicating that about one third of LAG-1 accumulation is promoted by GLP-1 transcriptional targets *lst-1* and *sygl-1*.

GLD-1 and GLD-2 repress LAG-1 accumulation in the proximal part of the progenitor zone (**Fig 3B; S3 Fig**). We found that in the *gld-2 gld-1* double null mutant, LAG-1 level was elevated ∼4.5-fold, as assessed at 25 cd from the distal tip. GLD-1 alone accounts for more than half of this repression, as LAG-1 level is elevated almost 3-fold in the *gld-1* single null mutant, which is consistent with *lag-1* mRNA being identified as a GLD-1 target in RNA pull-down experiments [44, 45]. As loss of *gld-2* function alone does not affect LAG-1 level, the remaining repression is apparently through the combined action of GLD-1 and GLD-2. The above results indicate that the fall in LAG-1 level in the proximal part of the progenitor zone is largely through translational repression by GLD-1 and GLD-2, as well as the absence of *lst-1* and *sygl-1* promoting LAG-1 accumulation, which are spatially restricted to the distal most ∼5 – 10 cd. Peak LAG-1 accumulation is also repressed by the combined activities of GLD-1 and GLD-2 (**Fig 3; S3 Fig**). This modest repression is presumably because of lower *gld-1* and *gld-2* activity in the distal most 10-cell diameters [4].

### Genome-wide identification of GLP-1 Notch – LAG-1 CSL transcriptional targets: ChIP-seq

*lst-1* and *sygl-1*, which were identified in a small candidate gene RNAi screen, are GLP-1 transcriptional targets that are redundantly required for the germline stem cell fate [21,22,42] (**Introduction**). However, it is possible that there are additional primary transcriptional targets of GLP-1 signaling that remain to be identified. Here, we undertook an unbiased genome-wide approach to identify transcriptional targets of GLP-1 signaling, through intersection of genes identified as directly bound by LAG-1, from <u>Ch</u>romatin immuno<u>p</u>recipitation (ChIP) experiments, with genes that are identified as requiring GLP-1 signaling for RNA accumulation, from RNA-seq transcriptomics analysis. LAG-1 CSL is the DNA binding protein that mediates transcription of GLP-1 and LIN-12 signaling targets, and was used in the ChIP-seq experiments. We first performed conventional ChIP-seq on whole mid-L4 worms, using the endogenous *lag-1* gene tagged with GFP and 3xFLAG, *lag-1(ar611[lag-1::GFP::3xFLAG])* (**S4 Fig**; gift from Iva Greenwald). We separately performed anti-FLAG or anti-GFP ChIP experiments; both antibodies can efficiently pulled-down tagged LAG-1 and gave a significant fold enrichment of *lst-1* and *sygl-1* DNA fragments compared to control genes in ChIP-qPCR (**S4C Fig, Materials and Methods**). Following high throughput sequencing, peaks were identified with greater than a two-fold enrichment compared to input control, and a false discovery rate (FDR <0.05). The Homer suite [46] was used to annotate the peaks to their nearest transcription start site (TSS). Seventy-five genes were identified as binding to LAG-1 from the intersection of the FLAG-IP (one biological replicate) and GFP-IP (two biological replicates) ChIP-seq (**S4 Fig; S3 Table**).

Known somatic LIN-12 signaling targets involved in vulval development, *lst-1* and *mir-61/250* [24, 47], and germline target *sygl-1* [21] were identified with each having a single major ChIP-seq peak that covers multiple canonical LAG-1/CSL binding sites. The *lag-1* gene contained multiple peaks, particularly in the large first intron that contains multiple canonical LAG-1/CSL binding sites. The *lag-1* first intron is also bound by a large number of transcription factors (∼50% of those analyzed to date, http://www.modencode.org/), but this is less than the definition of a Highly Occupied Target (HOT), where >65% of transcription factors bind to a region [48, 49]. Thus, the *lag-1* gene may function in a LIN-12 (and possibly GLP-1) signaling dependent positive autoregulatory feedback loop with LAG-1 in the soma, consistent with *lag-1* reporter gene analysis in the anchor cell – ventral uterine cell decision [50] (K. Luo and I. Greenwald, personal communication), although this does not appear to be the case in the germline (see above). We did not identify peaks for other reported LIN-12 or GLP-1 signaling somatic targets, such as *ref-1* in the embryo or *lip-1* in vulval development [51, 52]. This maybe a result of our analysis being from only a single stage (mid-L4) or because only a small number of cells are expressing the target gene under LIN-12 or GLP-1 control, resulting in only a small amount of total LAG-1 bound DNA, which is below the limit of detection in the ChIP-seq assay. We performed *de novo* discovery of over-represented DNA sequence motifs among the 75 genes and found the highest hit to be an 9-mer that contains the canonical LAG-1/CSL binding site (*p-value: e-^42^*)(**S4H Fig**). Thus, we believe that the bulk of the genes identified are bound by LAG-1 *in vivo*.

To identify genes whose transcription promotes the stem cell fate through direct LAG-1 binding, we performed germline specific LAG-1 ChIP-seq, where the approach was guided by our expression and germline autonomy analysis above. LAG-1 is modestly expressed in germline stem cells compared to much higher expression in late stage pachytene and diplotene oogenic germ cells and somatic cells (**Fig 2, 3**). This necessitated performing germline LAG-1 ChIP-seq on L4 stage hermaphrodites that lack oogenic germ cells in pachytene and diplotene. We generated a transgenic strain with the following components: (1) A fosmid transgene where the BioTag, a 23-amino acid peptide that is recognized and biotinylated by the *E. coli* enzyme BirA biotin ligase [53], was placed at the C-terminus of LAG-1 (*ozIs43[lag-1::3xFLAG::BioTag*, hereafter called *lag-1::BioTag*) (**S5A Fig; Materials and Methods**); (2) A transgene with germline specific expression of *E. coli BirA* biotin ligase from the *pie-1* promoter (*ckSi11[pie-1p::BirA::gfp]*); (3) The *lag-1* deletion allele, *tm3052*, which demonstrated that *lag-1::BioTag* in *ozIs43* produced functional, rescuing, LAG-1. Germline specific expression of BirA results in germline restricted biotinylation of LAG-1::BioTag, which can be pulled down by streptavidin beads (**Fig 4A**). However, because of high levels of endogenous *C. elegans* biotinylated proteins [54, 55] we were not able to directly pull-down the low levels of biotinylated LAG-1::BioTag. To overcome this issue, we performed sequential ChIP, first with anti-FLAG and then with streptavidin beads, followed by library construction directly on the beads due to ultra-low quantities of DNA following sequential ChIP (**Fig 4B**). Germline specific LAG-1 ChIP-seq data was analyzed as described for whole worm ChIP-seq.

**Figure 4.**
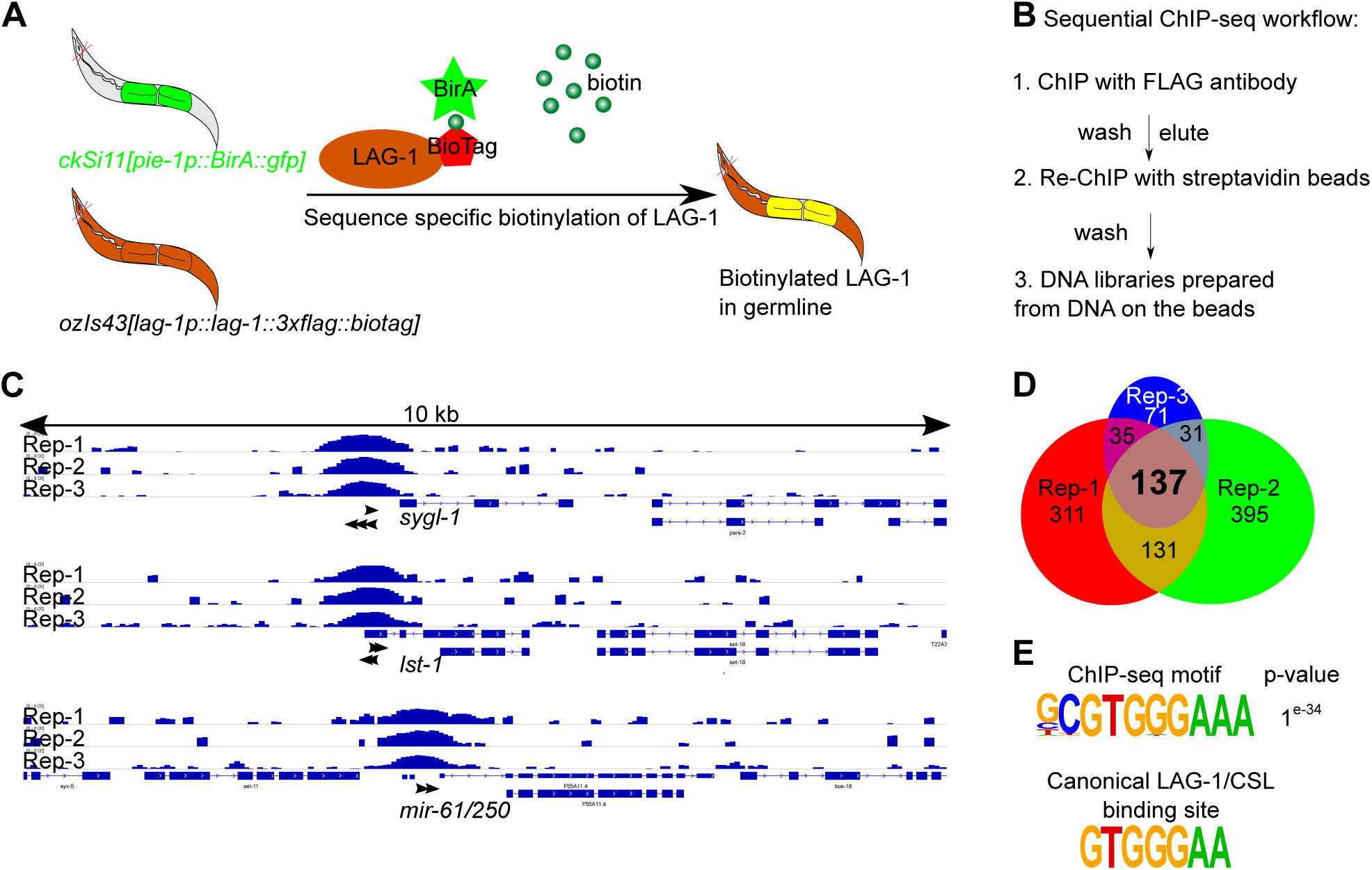
Genome-wide identification of germline-specific LAG-1 targets by sequential ChIP-seq. (A) Schematic showing germline specifically expressed BirA (in green) biotinylating LAG-1 containing the BioTag. The strain harbored two transgenes, i) *ckSi11[pie-1p::BirA::gfp],* where biotin ligase BirA was driven by germline specific promoter *pie-1,* and ii) *ozIs43[lag-1p::lag-1::3xFLAG::BioTag]*, see **S1 Table.** (B) The workflow of sequential ChIP-seq. Direct ChIP was not feasible due to the interference by high levels of endogeneous biotinylated proteins [54, 55] Sequential ChIP-seq was developed to overcome this issue and to facilitate DNA library preparation from ultra-low amount of DNA. (C) Genome browser tracks for *sygl-1, lst-1* and *mir-61/250* after sequential ChIP-seq. The raw reads were normalized to the control, and the signals are presented as log2-fold change after normalization. Black arrows, canonical LAG-1/CSL binding motif GTGGGAA [17,19,20]. (D) Venn diagram showing 137 genes were overlapped from three biological replicates of germline LAG-1 ChIP-seq analyses. Rep, replicate. (E) The over-represented motif discovered by HOMER suite with germline specific ChIP-seq data (top) and the reported canonical LAG-1/CSL binding motif (bottom) [17,19,20]. See **S4 Table.**

One hundred and thirty seven genes with germline specific LAG-1 peaks were identified in common between three biological replicates, including *lst-1*, *sygl-1* and *mir-61/250* (**Fig 4C-D**), providing biochemical support that *lst-1* and *sygl-1* are germline GLP-1 transcriptional targets. The *lag-1* gene was also found to contain multiple peaks. However, other linked genes on the fosmid (e.g., *zen-4*) and the selectable marker used (*unc-119*) also contained multiple peaks, which were absent in the whole worm ChIP-seq. Thus, they likely represent artificially peaks due to multiple integrated copies of *ozIs43*, therefore not considered to be specifically bound by germline LAG-1. *De novo* motif identification recovered the same 9-mer that contains the canonical LAG-1/CSL binding site (*p-value: e-^34^*) as found in the whole worm experiment, consistent with LAG-1 binding many of these genes *in vivo*, in germ cells. Thirty-six genes were identified in common between germline specific and whole worm that show LAG-1 ChIP-seq peaks, further supporting that LAG-1 binds to these genes *in vivo* (**S5E Fig**). Thus, the ChIP-seq analysis uncovered many genes bound by LAG-1 in germline, whether GLP-1 signaling regulates these genes is not clear.

### Genome-wide identification of GLP-1 Notch – LAG-1 CSL transcriptional targets: RNA-seq

To identify genes that require GLP-1 signaling for expression we performed RNA-seq comparing two strains, one with GLP-1 signaling ON, using *glp-1* <u>g</u>ain of function (gf) allele *ar202* that produces a large number of proliferating germ cells undergoing GLP-1 signaling at the restrictive temperature [15], and the other with GLP-1 signaling OFF, using the null allele *q175*. As described above, we employed the meiotic entry defective *gld-2 gld-1* double null mutant to allow examination of the effect of *glp-1* null in proliferating germ cells; *glp-1* gf was also placed in this background so that genotype was identical, except for *glp-1* activity status (**Fig 5A; Materials and Methods**). Further, we used dissected gonad preparations to significantly enrich for expression changes that occur in the germline. *In situ* hybridization and qRT-PCR was used to confirm that in GLP-1 ON, transcriptional target *lst-1* and *sygl-1* were expressed throughout the germline at significantly elevated levels, compared to GLP-1 OFF where expression was similar to background (**Fig 5B-C**, **S6A Fig**). RNA-seq was performed on 5 biological replicates, following ribosomal RNA removal, from GLP-1 ON and GLP-1 OFF dissected gonad RNA preparations. Heatmap and <u>p</u>rinciple <u>c</u>omponent <u>a</u>nalysis (PCA) demonstrated significant differences between the GLP-1 ON and GLP-1 OFF RNA-seq results (**S6B-C Fig**). We identified 94 genes whose RNA accumulation was dependent on GLP-1 signaling, with greater than 2-fold elevation of reads in GLP-1 ON versus GLP-1 OFF, FDR<0.05, and requiring greater than two counts per million reads (CPM) (**S5 Table; Materials and Methods;**).

**Figure 5.**
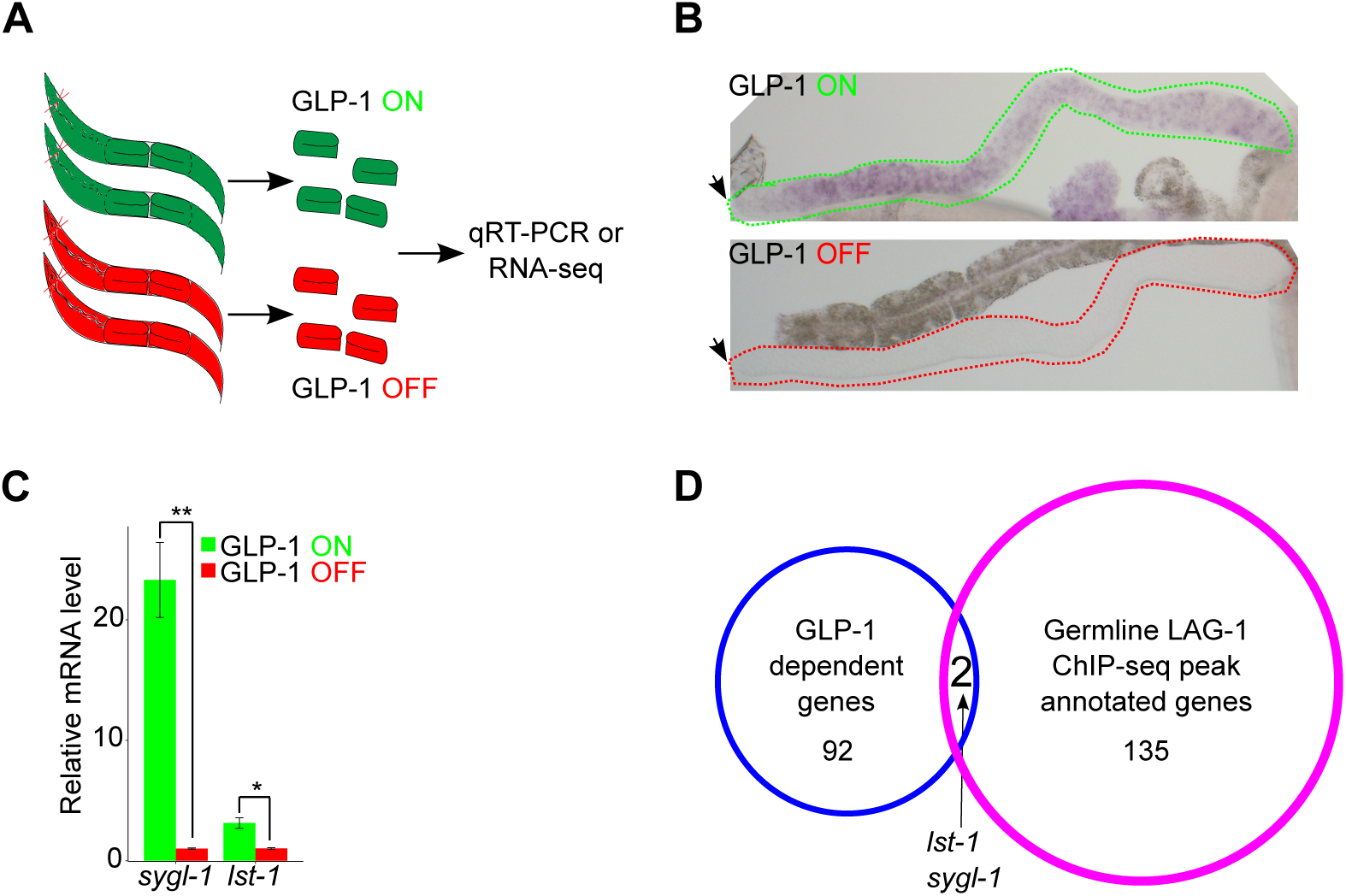
Genome-wide identification of GLP-1-dependent genes by transcriptomic analysis. (A) Schematic showing harvesting of dissected gonads, with tumorous germlines, for either qRT-PCR analysis or transcriptomic analysis. The genotype for **GLP-1 ON** animal (green) is *gld-2(q497) gld-1(q485); glp-1(ar202)* and **GLP-1 OFF** (red) animal is *gld-2(q497) gld-1(q485); glp-1(q175)*. (B) *In situ* hybridization (ISH) used to detect *sygl-1* mRNA expression in GLP-1 ON and GLP-1 OFF young adult animals. Dotted lines showing boundary of the gonad. Black arrow indicates distal end of the germline. (C) *sygl-1* and *lst-1* transcript levels analysis via qRT-PCR. The expression level for each gene in the GLP-1 OFF background was set as one. Three biological replicates were conducted and two tailed t-test was used for statistical analysis. P-value ≤ 0.01 (*); ≤ 0.001 (**). (D) Venn diagram identifying GLP-1 transcriptional targets through integrative genome-wide approach. Five biological replicates were used to conduct transcriptomic analysis and identified 94 GLP-1-dependent genes (blue circle). Two GLP-1 transcriptional targets were defined as genes whose mRNA expression was controlled by GLP-1 and also had LAG-1 bound to their promoter regions (red circle, data from Fig 4D). See **S5 Table**.

To identify primary GLP-1 signaling germline transcriptional targets we intersected the 137 genes identified as bound by LAG-1 with the 94 genes whose RNA accumulation was dependent on GLP-1 signaling, which yielded only two genes, *lst-1* and *sygl-1* (**Fig 4, 5**). Three additional genes have been reported as germline GLP-1 signaling targets, *fbf-2*, *utx-1*, and *lip-1* [30–32]. We did not observe LAG-1 germline ChIP-seq peaks (**S5B Fig**) for *fbf-2*, *utx-1* or *lip-1*. For GLP-1 dependent RNA accumulation in GLP-1 ON versus OFF, *fbf-2* mRNA level was unchanged, *utx-1* mRNA level was below the 2 CPM cutoff, while *lip-1* mRNA increased 2.3 fold in GLP-1 ON. Consistent with unchanged mRNA level, an *FBF-2::fbf-2* 3’UTR reporter driven by the heterologous *pie-1* promoter gives a qualitatively similar FBF-2 progenitor zone protein accumulation as observed in wild type [30, 56] (see below), supporting GLP-1 signaling independent control of *fbf-2* mRNA accumulation. Prior work indicated that *lip-1* mRNA accumulation in the progenitor zone was principally controlled by FBF-1 mediated degradation [31], inconsistent with it being a primary GLP-1 target. The RNA-seq method we employed does not recover RNAs less than ∼100nt. Thus, it remains possible that small RNAs, including *mir-61/250*, are transcriptional targets of GLP-1 signaling (**Fig 4C-D**). Together, the above experiments provide robust *in vivo* biochemical support for *lst-1* and *sygl-1* being mRNA transcriptional targets of germline GLP-1 signaling, and potentially being the only primary GLP-1 signaling transcriptional targets.

### FBF-2 accumulation is controlled by GLP-1 signaling transcriptional targets *lst-1* and *sygl-1*

Previous work showed that FBF-2 accumulation is enriched in the progenitor zone, with peak FBF-2 accumulation dependent on GLP-1 signaling [30]. We used CRISPR/Cas9 tagged *fbf-2(q932*[*3xV5::fbf-2*]) [23], hereafter called *fbf-2::V5*, to quantitatively examine FBF-2 accumulation, following anti-V5 antibody staining in dissected gonads from young adult hermaphrodites (**Materials and Methods**). We found that FBF-2::V5 displayed a peak of accumulation at 8 – 13 cd from the distal tip, followed by a somewhat gradual fall to low proximal levels by ∼35 cd (**Fig 6A-B**), similar to previously reported [30], with peak accumulation ∼4 fold higher than base. In *gld-2 gld-1* double null mutant germlines, the FBF-2::V5 peak was ∼80% of wild type, with a the fall more rapid than in wild type and a flat base from ∼18 cd through 35 cd, with the peak also ∼4 fold higher than base (**Fig 6A, C**). In the *gld-2 gld-1; glp-1* triple null mutant germlines, FBF-2::V5 levels are low throughout the progenitor zone (**Fig 6A, C-D**), with the level ∼4 fold lower than in the *gld-2 gld-1* double mutant. Thus, peak FBF-2 accumulation requires GLP-1 signaling, with ∼15% of FBF-2 accumulation being GLP-1 signaling independent, as previously reported [30]. Given that FBF-2 accumulation appears to be controlled post-transcriptionally, and the *lst-1* and *sygl-1* promote peak LAG-1 accumulation in the progenitor zone [56] (**Fig 3**), we next examined whether LST-1 and SYGL-1 promoted peak FBF-2 accumulation. Analysis of *gld-2 gld-1 lst-1 sygl-1* quadruple null mutant germlines showed low FBF-2 accumulation throughout the progenitor zone (**Fig 6**), consistent with LST-1 and SYGL-1 being required for peak FBF-2 accumulation. In the proximal progenitor zone, GLD-1 appears to promote FBF-2 accumulation as the level is lower in the *gld-1* null mutant (**S7 Fig**). Since GLD-1 acts in translational repression, the effect on FBF-2 accumulation is presumably indirect. In summary, FBF-2 is a secondary protein accumulation target of GLP-1 signaling as *glp-1* dependent peak FBF-2 progenitor zone levels appear to be explained through post-transcriptional regulation by *lst-1* and *sygl-1* activity (**Fig 6E**).

**Figure 6.**
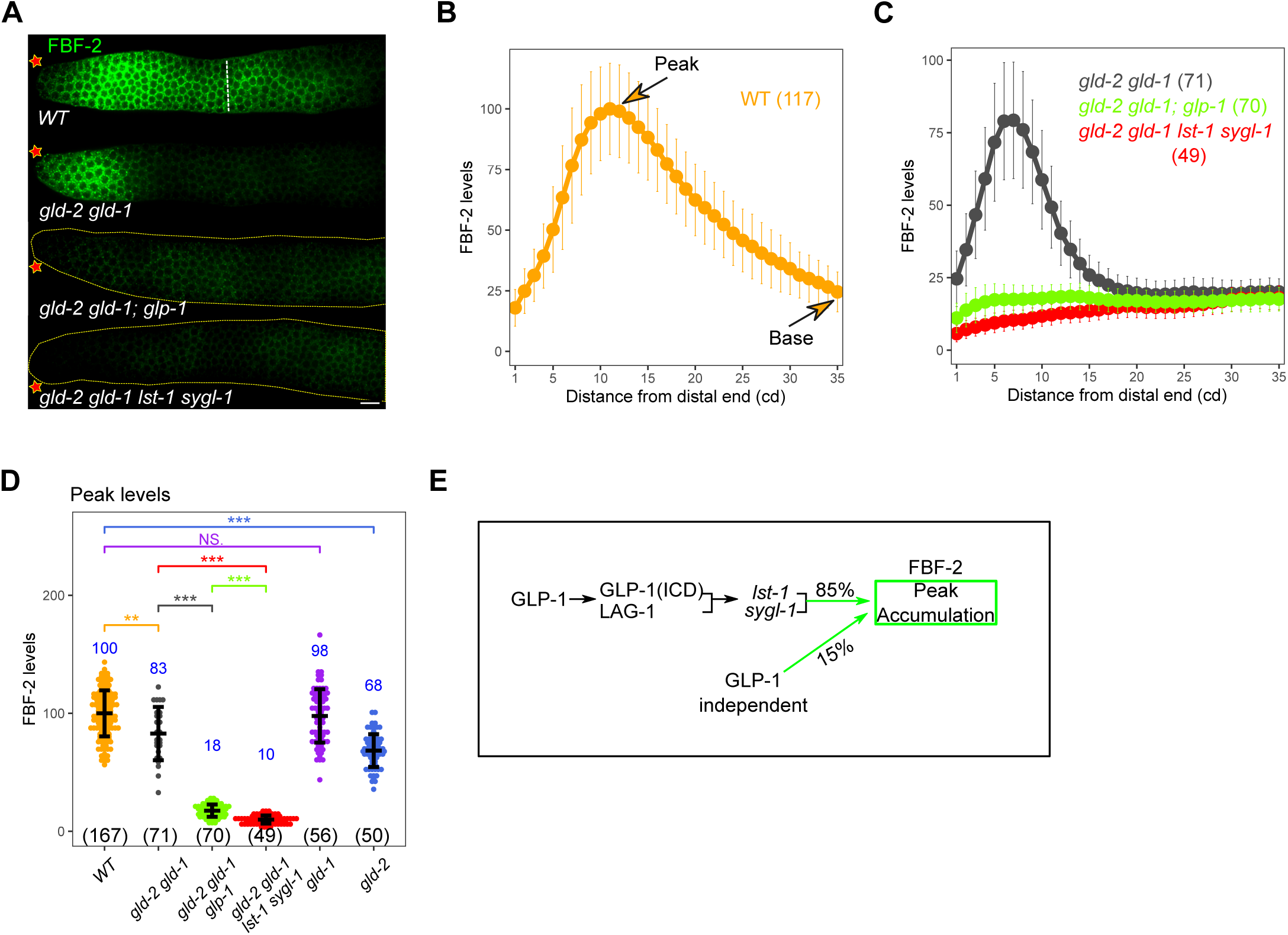
FBF-2 accumulation is post-transcriptionally controlled by GLP-1 transcriptional targets LST-1 and SYGL-1. (A) Images of V5-stained (FBF-2, green, from *fbf-2(q932[3xV5::fbf-2])* germlines from dissected young adult hermaphrodites of the indicated genotype. See **S1 Table** for the complete genotypes. Asterisk, distal end; dashed white line, meiotic entry. Scale bar is 10 μm. (B - D) Plot of FBF-2 levels (B - C) and comparison of peak FBF-2 accumulation (D) for indicated genotype. Numbers indicate mean values of FBF-2 level for each genotype (in blue) and numbers in bracket shows the sample size. Dots, mean (B-C) or data points (D); Error bars, mean ± SD. P-value ≤ 0.01 (*); ≤ 0.001 (**); ≤ 0.0001 (***); > 0.01 non-significant (NS.). (E) Model depicting genetic control of FBF-2 peak levels and percent contribution to peak FBF-2 accumulation. Total of 85% of FBF-2 peak level can be attributed to GLP-1 signaling through GLP-1 transcriptional targets *lst-1* and *sygl-1*.

### Germline LAG-1 functions in transcriptional activation of *lst-1* and *sygl-1*

Only two of 137 germline LAG-1 ChIP-seq peaks are associated with GLP-1 dependent activation of target transcription. The remaining peaks may reflect LAG-1 acting independent of GLP-1 signaling, either as a transcriptional activator or repressor. In Drosophila, it is known that in the absence of Notch ICD, *Su(H)* CSL functions in actively repressing Notch transcriptional target genes [57]. To test if LAG-1 has a GLP-1 signaling independent transcriptional function in the germline, we performed LAG-1 AID (see above) followed by gonad dissection and RNA-seq (**Materials and Methods**; **Fig 2D, Fig 7**). We used a strain containing *lag-1::degron::HA*, *glp-1(ar202)* gf that at the restrictive temperature will have the bulk of germ cells undergoing GLP-1 signaling, and the meiotic entry defective double mutant *gld-2 gld-1*, to allow analysis of proliferating germ cells, with or without auxin treatment.

**Figure 7.**
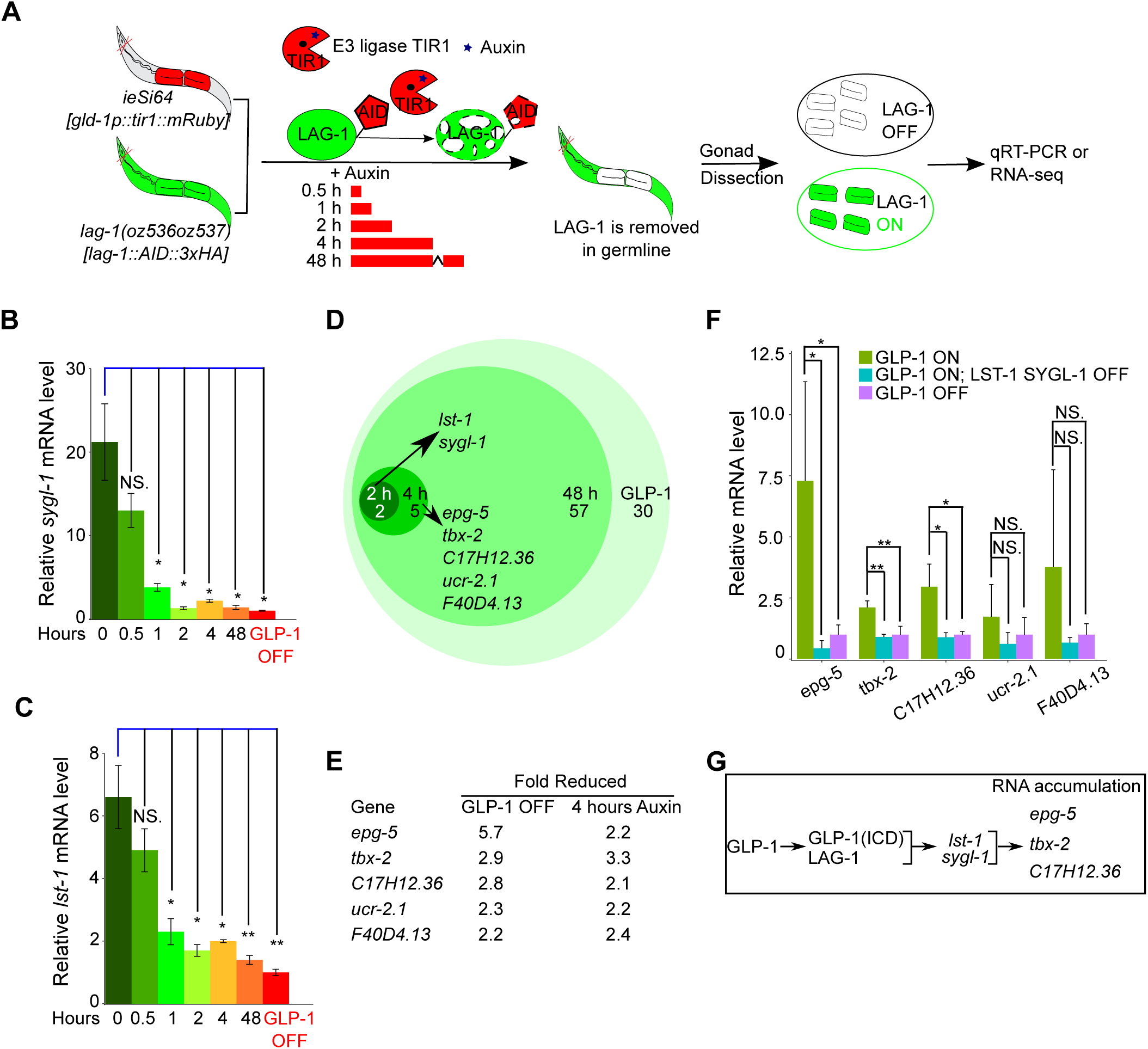
Time-course transcriptomic analysis to identify LAG-1-dependent genes in the germline. (A) Schematic showing time-course qRT-PCR and transcriptomic analysis following germline-specific degradation of LAG-1. This strain harbors germline-expressed TIR1, an ubqiutin E3 ligase that drives degron containing target protein degradation through proteolysis in the presence of co-factor auxin [37]. The CRISPR allele *lag-1(oz536oz537)* has degron fused to the C-terminus of LAG-1, which is recognized by TIR1, resulting in germline-specific degradation of LAG-1. The complete genotype for this strain is *gld-2(q497) gld-1(q485); glp-1(ar202); lag-1(oz536oz537[lag-1::degron::3xHA]); ieSi64[gld-1p::TIR1::mRuby::gld-1 3’UTR].* Isolated gonads, after dissection, are used to quantify the abundance of RNAs, either through qRT-PCR or transciptomic analysis. (B & C) Quantification of two GLP-1/LAG-1 targets *sygl-1* and *lst-1* mRNA abundance after auxin treatment at different time points. Three biological replicates were conducted and two-tailed t-test was used for statistical analysis. P-value ≤ 0.01 (*); ≤ 0.001 (**); ≤ 0.0001 (***); > 0.01 non-significant (NS.). (D) Venn diagram showing genes whose RNA expression is dependent on LAG-1 at the various time points through transcriptomic analysis. Genes observed at an earlier time are also dependent at later time point(s), but not shown for simplicity. The presented genes were also controlled by GLP-1. Four biological replicates were conducted for each time point. (E) Expression of 5 genes was reduced after 4 hrs auxin treatment, in addition to *lst-1* and *sgyl-1* (identified at 2 hrs auxin treatment). Shown are the genes’ relative fold reduction in both GLP-1 OFF and LAG-1 OFF (via 4 hrs auxin treatment) from transcriptomic analysis. (F) qRT-PCR analysis for genes expression level from (E) in different mutant backgrounds. The genotypes are, **GLP-1 ON**: *gld-2(q497) gld-1(q485); glp-1(ar202).* **GLP-1 OFF**: *gld-2(q497) gld-1(q485); glp-1(q175).* **GLP-1 ON LST-1 SYGL-1 OFF**: *gld-2(q497) gld-1(q485) lst-1(ok814) sygl-1(tm5040); glp-1(ar202).* Five biological replicates were conducted for each genotype and two-tailed t-test was used for statistical analysis. P-value ≤ 0.01 (*); ≤ 0.001 (**); ≤ 0.0001 (***); > 0.01 non-significant (NS.). (G) Model depicting genetic control of RNA accumulation for genes from (F). See **S6-7 Tables**.

We performed RNA-seq following 48 hrs of auxin treatment from the L1 stage at 25°C, a time where LAG-1 is undetectable, and the same time point where GLP-1 ON and OFF RNA-seq was performed (**Fig 5**). Heatmap and PCA demonstrated that changes in RNAs from LAG-1 ON (minus auxin) and LAG-1 OFF (plus auxin) were significantly different, while biological replicates were similar (**S8A-B; S6 Table**). Ninety-four genes were identified whose expression was dependent on LAG-1 (activated genes), where ∼70% (64) of these were also GLP-1 dependent genes (**S8C Fig; S5-6 Tables**). Only two of the LAG-1 activated genes overlapped with LAG-1 germline specific ChIP-seq peak containing genes, *lst-1* and *sygl-1* (**S8E Fig**). Forty-eight genes were identified that were LAG-1 repressed, where ∼80% of these were also GLP-1 repressed genes (**S8D Fig**). There was no overlap between LAG-1 repressed genes and germline LAG-1 ChIP-seq peak genes (**S8F Fig**), which is not consistent with LAG-1 functioning in transcriptional repression in the *C. elegans* germline. We note that the LAG-1 activated/repressed genes list and the GLP-1 signaling activated/repressed gene list derive from strains that functionally differ in at least two ways that may have resulted in incomplete overlap of gene lists. First, LAG-1 AID may not completely eliminate LAG-1 protein, while the *glp-1(q175)* allele is null. Second, while both use dissected gonads for RNA-seq, LAG-1 AID degrades LAG-1 specifically in the germline, while *glp-1(q175)* lacks GLP-1 signaling in both the germline and the soma. The above results indicate that while LAG-1 has an essential function in transcriptional activation of *lst-1* and *sygl-1* for germline GLP-1 signaling, it does not have an essential GLP-1 signaling independent function in either transcriptional activation or repression.

### Primary versus secondary GLP-1 signaling mRNA targets

The LAG-1 AID system described above provides a route to distinguish between genes whose expression is a primary or a secondary target of GLP-1 signaling/LAG-1 activity. By performing a time course of LAG-1 degradation, using RNA-seq as the readout, genes whose RNA levels are transcriptionally controlled by GLP-1 signaling should change expression earlier, while genes whose RNA level is indirectly controlled/a secondary effect of GLP-1 signaling, should change expression later.

We performed a LAG-1 AID time course, harvesting RNA following 0.5, 1, 2, 4 and 48 hrs of auxin treatment (**Fig 7**). *sygl-1* and *lst-1* mRNA levels, as assessed by qRT-PCR, dropped ∼3 fold by 1 hr of auxin treatment and by 2 hrs were not significantly different from 48 hrs of auxin treatment or GLP-1 OFF. Staining for LAG-1::degron::3xHA indicated that at 2 and 4 hrs of auxin treatment, LAG-1 was at a low/undetectable level, while staining for WAPL-1 show no change in the size of the progenitor zone (**S9 Fig**), indicating that the switch of all the stem cell to meiotic development, following loss of LAG-1, was only just beginning. RNA-seq was therefore performed at 2 and 4 hrs, time points, where there was minimal pleiotropy from loss of LAG-1. Multidimensional scaling (MDS) plots revealed that the 0, 2, and 4 hrs auxin treatment clustered together, but separately from 48 hrs auxin treatment, indicating that there are few changes in expression at early times following degradation of LAG-1 compared to strong loss of LAG-1 (**S8G Fig**). After 2 hrs of auxin treatment, among the GLP-1 dependent genes, only *lst-1* and *sygl-1* were identified as LAG-1 mRNAs targets (**Fig 7B-D**). By 4 hrs of auxin treatment, 5 additional genes were identified whose RNAs displayed both LAG-1 and GLP-1 dependent accumulation (**Fig 7D-E**). The new 5 genes from the 4 hrs time point could represent additional GLP-1 - LAG-1 transcriptional targets, which have longer RNA half-lives than *lst-1* and *sygl-1*, or they may represent changes in RNA level that are secondary effects of GLP-1 signaling. However, we did not detect germline LAG-1 ChIP-seq peaks or canonical LAG-1/CSL binding sites in these 5 genes, indicating that they do not fit the criteria of a primary GLP-1 signaling transcriptional target. From these results, and our previous finding that *lst-1* and *sygl-1* promote LAG-1 and FBF-2 protein accumulation, we reasoned that RNA accumulation for these 5 genes may also be dependent on *lst-1* and *sygl-1* activity.

We first examined the kinetics of loss of LST-1 and SYGL-1 following LAG-1 AID. Peak LST-1 falls ∼5 fold at 2 hrs and ∼10 fold at 4 hrs, while SYGL-1 falls ∼3 fold at 2 hrs and 10-fold at 4 hrs auxin treatment (**S10 Fig**). Given their rapid loss, the significant reduction of LST-1 and SYGL-1 could be responsible for the reduction in RNA level of the 5 genes at the 4 hrs auxin treatment time point. To test this, we measure RNA level by qRT-PCR for the 5 genes (*epg-5*, *tbx-2*, C17H12.36, *ucr-2.1*, F40D4.13) from dissected gonads with GLP-1 ON, GLP-1 ON but lacking *lst-1* and *sygl-1* activity (LST-1 SYGL-1 OFF) and GLP-1 OFF (see **Fig 7F** and legend for full genotype). For *epg-5*, *tbx-2* and C17H12.36, higher RNA accumulation in GLP-1 ON was dependent on *lst-1* and *sygl-1* activity. Thus, *epg-5*, *tbx-2* and C17H12.36 are indirect or secondary targets of GLP-1 signaling, with their RNA accumulation dependent on the downstream *lst-1* and *sygl-1* gene activity. It is likely that other genes whose RNA level are dependent on both LAG-1 at 48 hrs and GLP-1 signaling are secondary targets that also rely on *lst-1* and *sygl-1* activity. The LAG-1 AID time course provides an estimate of the maximum half-life of the *lst-1* and *sygl-1* mRNAs, 1 hr, and the LST-1 and SYGL-1 proteins, 2 hrs, at 25°C. Taken together, the LAG-1 AID time course results supports findings from the initial genome-wide analysis that *lst-1* and *sygl-1* are likely the only two mRNA transcriptional targets of GLP-1 signaling.

## Discussion

A central question for the *C. elegans* germline stem cell system is the identity of the GLP-1 Notch dependent transcriptional targets that promote the stem cell fate. We performed a genome-wide search for transcriptional targets through germline specific ChIP-seq analysis with LAG-1, the Notch signaling CSL DNA binding protein homolog, and intersected these results with transcriptomics analysis under conditions of the presence or absence of GLP-1 signaling. We also employed a time-course transcriptomics analysis following auxin inducible degradation of LAG-1 to distinguish between genes whose RNA level was a primary or secondary response of GLP-1 signaling. These approaches identified only *lst-1* and *sygl-1*, two previously described GLP-1 transcriptional targets, and supports the possibility that there are no additional direct mRNA transcriptional targets of germline GLP-1 signaling. We further report examples of GLP-1 dependent secondary targets, whose RNA level or protein accumulation was dependent on *lst-1* and *sygl-1*. We elaborate on these and other findings below.

### GLP-1 - LAG-1 transcriptional control of the germline stem cell fate

We present three lines of genome-wide molecular support that *lst-1* and *sygl-1* are direct GLP-1 signaling transcriptional targets. First, we identified robust germline LAG-1 ChIP-seq peaks in the promoters of *lst-1* and *sygl-1*, overlapping the position of consensus LAG-1/CSL binding sites. Second, transcriptomics analysis demonstrated GLP-1 signaling and germline LAG-1 dependent *lst-1* and *sygl-1* mRNA accumulation. Third, in a LAG-1 AID time course, *lst-1* and *sygl-1* mRNAs were the first RNAs to fall, as early as 0.5 to 1 hr after auxin treatment, indicating that their mRNA loss was a direct/primary effect of reduced LAG-1 activity. Combined with prior work that *lst-1* and *sygl-1* transcription in the distal progenitor zone is dependent on GLP-1 signaling and that distal *sygl-1* reporter gene expression requires canonical LAG-1/CSL binding sites [21, 22], our results strengthen the conclusion that *lst-1* and *sygl-1* are direct transcriptional targets.

This work supports the possibility that there may not be any additional direct mRNA transcriptional targets of GLP-1 signaling. Our results are not consistent with three reported genes, *fbf-2*, *lip-1* and *utx-1*, being GLP-1 transcriptional targets [30–32]. ChIP-seq peaks were not observed for the three genes. *fbf-2* mRNA level was unchanged with or without GLP-1 signaling or LAG-1 protein. For *lip-1*, while GLP-1 signaling and LAG-1 promoted mRNA accumulation, in the LAG-1 time course *lip-1* mRNA level was unchanged at 2 and 4 hrs of auxin treatment, but was reduced at 48 hrs (**S5-7 Tables**). The delayed fall in *lip-1* mRNA level is consistent with an indirect/secondary effect of GLP-1 signaling and LAG-1 activity. For *utx-1*, its mRNA read count was below the 2 CPM cut-off for reliable RNA level assessment from transcriptomics analysis. While we identified genes whose RNA accumulation was GLP-1 signaling dependent (92 genes) or germline LAG-1 dependent (92 genes), other than *lst-1* and *sygl-1*, they lacked germline or whole worm LAG-1 ChIP-seq peaks. From the LAG-1 AID time course, five genes were identified where their RNA level was dependent of LAG-1 at 4 hrs of auxin treatment, raising the possibility that they were direct/primary targets of LAG-1 activity. However, in addition to the five genes lacking LAG-1 ChIP-seq peaks, RNA accumulation for three of the genes was dependent on *lst-1* and *sygl-1*. Thus, our genome-wide studies did not identify any new candidate GLP-1 signaling mRNA transcriptional target genes. However, the RNA-seq approach employed cannot assess the level of small RNAs (less than ∼100nt), and a number of small RNA genes were identified as containing germline LAG-1 ChIP-seq peaks (e.g, *mir-61/250*, five 21 U RNA genes). Thus, it is possible that there are small RNA genes that are germline GLP-1 signaling transcriptional targets. We note that because of the low amount of DNA that was obtained for the germline LAG-1 ChIP-seq experiments, it is possible that we missed some weak ChIP-seq peaks. We also note that all of the transcriptomics analyses were performed using dissected gonads from the *gld-2 gld-1* meiotic entry double mutant background, to allow germline RNA level analysis in the absence of *glp-1* or *lag-1* activity. We cannot rule out the possibility that the absence of *gld-1* and *gld-2* activity may mask GLP-1 signaling dependent changes in RNA levels. Nevertheless, the observation that the premature meiotic entry phenotype of the *lst-1 sygl-1* null double mutant is the same as *glp-1* null [21] indicates that no additional transcriptional targets genes are necessary to account for germline GLP-1 signaling.

### LAG-1 expression and function

We found that *lag-1* is germline autonomously required for the stem cells fate, as predicted from our current understanding of Notch signaling and notwithstanding strong LAG-1 accumulation in all somatic gonad cells. In the soma, LAG-1 appears to function in a positive autoregulatory feedback loop to promote accumulation in cells undergoing the LIN-12 dependent AC/VU decision [50] (K. Luo and I. Greenwald, personal communication). Consistent with this possibility, we found multiple LAG-1 ChIP-seq peaks in the *lag-1* gene in the whole worm ChIP-seq experiments. In the germline, however, *lag-1* mRNA was found at a constant level in the progenitor zone, in the presence or absence of *glp-1* activity. These findings are consistent with germline *lag-1* transcription being independent of GLP-1 signaling, with a *lag-1* positive transcriptional feedback loop not active in the germline. We note that the Notch signaling dynamics in the *C. elegans* germline differs from well-known lateral signaling examples, which may then affect mechanistic aspects of transcriptional control. GLP-1 signaling and *lst-1* and *sygl-1* activity are continuously required in the *C. elegans* germline to promote the stem cell fate, from the L1 larval stage through at least mid-adulthood, under optimal growth conditions [4]. Thus, with the possible exception of the early L1 stage, germline stem cells are born undergoing GLP-1 signaling, and only lose GLP-1 signaling when cells are displaced away from the DTC niche (ON>OFF). In contrast, cells undergoing lateral signaling initially lack Notch signaling, then undergo Notch signaling, and then may continue or downregulate Notch signaling (OFF>ON>OFF). Such differences in Notch signaling dynamics are likely mirrored in the transcriptional states of respective cells and may provide an explanation for differences observed between the *C. elegans* germline, somatic cells, and other organisms in Notch mediated transcriptional control, and may be a reason for the absence of the LAG-1 positive autoregulatory transcriptional feedback loop in the germline.

Mammalian CBF1 and Drosophila Su(H), orthologs of LAG-1, can act as transcriptional repressors in the absence of Notch signaling [57, 58]. Following germline specific loss of LAG-1, we found that *lst-1* and *sygl-1* RNA levels dropped significantly, equivalent to the absence of GLP-1 signaling. Thus, LAG-1 is required for GLP-1 dependent expression and, correspondingly, does not appear to function in repression of transcriptional targets *lst-1* and *sygl-1*. In the genome-wide studies, none of the germline LAG-1 repressed genes (RNA level elevated following LAG-1 AID) contain germline LAG-1 ChIP-seq peaks, indicating that transcriptional repression is not a general property of LAG-1 in the germline.

We identified 137 genes with LAG-1 ChIP-seq peaks but only two of these genes are transcriptionally controlled by GLP-1 signaling/LAG-1. This disconnect between the significantly larger number of genes that are bound by a transcription factor compared to the smaller number of genes that are regulated by the transcription factor is similarly observed in yeast and mammalian cells [59, 60]. The biological significance of this disconnect is currently unclear.

We found that LAG-1 protein accumulation is spatially restricted, high in the distal most 5 cd (peak) from the tip and then falls 7-fold to a base level ∼17 cd from the tip. Importantly, the germ cells with peak LAG-1 accumulation correspond to those where GLP-1 dependent nascent *lst-1* and *sygl-1* transcripts are observed [22, 42]. While GLP-1 signaling is not controlling *lag-1* mRNA level, we nevertheless found that 61% of peak LAG-1 accumulation was GLP-1 signaling dependent, suggesting a posttranscriptional mechanism (**Fig 3E**). This is consistent with the observation that many genes in the distal germline are regulated posttranscriptionally through the 3’UTR [61, 62]. We found that ∼50% of the GLP-1 dependent LAG-1 accumulation required *lst-1* and *sygl-1* activity. The basis for the remaining ∼50% of GLP-1 dependent LAG-1 accumulation is not known. Perhaps GLP-1(ICD) is stabilizing LAG-1 protein. The fall in LAG-1 levels in the proximal progenitor zone can be attributed, at least in part, to GLD-1 and GLD-2 activity. We speculate that the spatial restriction of LAG-1 contributes to controlling the size of the stem cells pool, with peak levels required for efficient *lst-1* and *sygl-1* transcription in the first 5 cd, and that lower levels in the proximal progenitor zone decrease the probability of stochastic GLP-1 signaling triggering transcription of *lst-1* and *sygl-1* in cells as they progress toward meiotic development.

### GLP-1 signaling indirectly mediates control of RNA and protein levels through LST-1 and SYGL-1

We found a number of examples of GLP-1 dependent RNA and protein accumulation that are through the activity of *lst-1* and *sygl-1*. Lamont et al. (2004) reported that distal peak FBF-2 accumulation requires GLP-1 signaling. We found that the *glp-1* dependent peak FBF-2 accumulation can be explained by the activity of *lst-1* and *sygl-1* (**Fig 6**). FBF-2 activity functions in repression of the GLD-1 and GLD-2 meiotic entry pathways. Additionally, as described above, ∼60% of peak LAG-1 accumulation is GLP-1 dependent, about 50% of which can be attributed to *lst-1* and *sygl-1* activity. From the LAG-1 AID time course, we identified 5 genes whose RNA accumulation depended on LAG-1, as well as *glp-1* activity. For *epg-5*, *tbx-2* and C17H12.36 we found that their GLP-1 and LAG-1 dependent expression can be explained by the requirement for *lst-1* and *sygl-1*activity. The remaining two genes, *ucr-2.1* & F40D4.13, show a trend to *lst-1* and *sygl-1* dependence, although it is not statistical significant with the number of replicates examined. Previous work reported that GLP-1 signaling inhibits GLD-1 accumulation in the distal progenitor zone [40] and this posttranscriptional repression required *lst-1* and *sygl-1* activity [23]. Thus, the bulk of germline gene expression changes ascribed to GLP-1 signaling, and *lag-1* function, can be attributed to the activity of transcriptional targets *lst-1* and *sygl-1*. These findings are consistent with the possibility that there are no additional mRNA targets of GLP-1 signaling that promote the stem cell fate.

LST-1 and SYGL-1 have been reported to function in conjunction with FBF-1 and FBF-2 in direct translational repression/mRNA destabilization in the posttranscriptional inhibition of GLD-1 accumulation [23, 28]. FBF-1 has also been shown to function in translational repression/mRNA destabilization in the posttranscriptional inhibition of FBF-2 accumulation [30]. In contrast, we find that *lst-1* and *sygl-1* function in posttranscriptional activation of FBF-2 and LAG-1 accumulation. Future work will be necessary to determine if *lst-1* and *sygl-1* act with or separately from FBF-1 and FBF-2 in posttranscriptional activation of FBF-2 and LAG-1 accumulation. Similarly, *lst-1* and *sygl-1* promote *epg-5*, *tbx-2* and C17H12.36 RNA accumulation, and the mechanism by which this occurs remains to be determined.

## Materials and Methods

### Strain maintenance

Unless otherwise noted, *C. elegans* strains were maintained at 20°C through conventional methods [63]. The animals were grown on NGM plates seeded with OP50 bacteria. *glp-1(ar202)* is a temperature <u>s</u>ensitive (*ts*) allele and strains with this allele were maintained at 15°C. Strains with complex genotypes were constructed by standard genetic recombination and segregation methods and where necessary, balancer chromosomes were employed to maintain sterile mutations. A complete list of strains used in this study is provided in **S1 Table**.

### Generation of *lag-1* CRIPSR alleles and transgenes expressing *lag-1* and BirA

*lag-1(oz530[lag-1::3xHA])* (**Fig 2A**) was generated through *in vitro* assembled RNA protein complex (RNP) CRISPR [64, 65]. Briefly, each component was pooled in volume of 20 µl with final concentration as follows before injection: 0.25 ng/µl HiFi Cas-9 (IDT, #1081061), 30 µM tracrRNA (IDT, #1072534), 15 µM *lag-1* specific crRNA(atacagtaatcccgcgagag<u>NGG</u>) (IDT, Alt-R™), 0.02 µM *lag-1* specific single strand DNA repair templates (3xHA sequences are underlined) (cctacaaatcattggaacgacatggaccgtgcagaattgtgtccaattac<u>TACCCTTACGACGTGCCAGATTACGCTTACCCCTACGACGTACCAGACTACGCCTACCCATACGACGTCCCAGACTACGCTTAG</u>attAAactctcgcgggattactgtatctttatattgtctcctaatttctcccaattcgt) (IDT), 15 µM *pha-1* crRNA and 0.02 µM *pha-1(e2123)* specific repair template [66]. The injected animals were raised at 25°C to identify *pha-1(e2123)* rescued animals. PCR was used to screen for edits (see **S2 Table** for oligonucleotide information). Generated alleles were then examined by Sanger sequencing to ensure there were no extraneous mutations. One allele, *lag-1(oz530),* outcrossed twice with wild type, was used for further analysis.

*lag-1(oz536oz537[lag-1::degron::3xHA])* was used to assess LAG-1 protein function in the germ cells (**Fig 2A**). This allele was generated through the Self Excising Cassette (SEC) method [67], due to insert size. The SEC allele contains the *sqt-1* roller marker and hygromycin antibiotic selection marker to facilitate screening [67]. Briefly, each component was mixed in a final volume of 20 µl with the following concentrations prior to injection of wild type animals: 50 ng/µl Cas9 plasmid (pDD121, *Cas9* driven by *eft-3* promoter), 50 ng/µl *lag-1* small guide RNA (sgRNA) plasmid (*lag-1* sgRNA sequence <u>atacagtaatcccgcgagag</u> was cloned into plasmid DR274 U6 through *BsaI* site), 10 ng/µl *lag-1* specific repair template and 2.5 ng/µl *myo-2p::gfp* co-injection marker. The *lag-1* specific repair template was constructed through Golden Gate cloning method [68]. The injected animals were raised at 20°C for three days prior of adding 500 µl of 5 mg/µl hygromycin. Six days later, animals that survived the antibiotic treatment and without *myo-2p::gfp* injection marker were screened for inserts by PCR (see **S2 Table** for oligonucleotide information). *lag-1(oz536)*(rollers, with SEC in) was verified through Sanger sequencing, followed by two times outcrosses with wild type prior to heat shock to remove the SEC [67] to generate *lag-1(oz536oz537)*

*lag-1(ozIs43[lag-1::3xFLAG::BioTag + unc-119])* fosmid transgenic line (**S5A Fig**) was generated through insertion of the 3xFLAG and BioTag encoded peptide sequence at the C-termimus of LAG-1 in pCC1FOS vector (Source BioScience, CBGtg9050C1288D) by Recombinering [69]. BioTag is a 23 amino acid peptide that can be recognized and specifically biotinylated by biotin ligase BirA [54]. All transgenic alleles were generated through bombardment and *unc-119* rescue; three alleles were found to rescue the phenotype of *lag-1(tm3052)*. One of the alleles, *lag-1(ozIs43)* was chosen to generate strain BS1193 (**S1 Table**) and used for germline LAG-1 ChIP-seq analysis (**Fig 4**).

*ckSi11[pie-1p::BirA::gfp]*. *C. elegans* codon-optimized BirA was synthesized with artificial introns (Genescript) and tagged at the C-terminus with GFP. Codon optimization was necessary for efficient germline BirA expression. This construct was cloned with the *pie-1* promoter and the *npp-9* 3’ UTR to promote germline expression and incorporated into the chromosome I safe-harbor site *ttTi4248* using MosSCI [70]. GFP expression was observed in the L4 and adult germline, deposited into embryos, and visible until at least the 250 cell stage.

### Auxin treatment

The auxin treatment was used to degrade degron tagged LAG-1 in the germline. Indole 3-acetic acid (IAA), the native plant hormone used in this study was ordered from Alfa Aesar (#A10556) [37]. Like Zhang et al. 2015, we noticed that high concentration of auxin (4 mM) inhibited OP50 bacteria growth. Therefore, all treatments were done in NGM plates supplemented with 1 mM auxin (auxin plates). The un-seeded auxin plates were stored at 4°C in the dark, and used within two weeks. The auxin plates were seeded with OP50 bacteria and kept at 20°C for 24 hours before use. The auxin treatment was conducted either at 20°C (**Fig 2E**) or at 25°C (**Fig 7**, **S8-10 Fig**) with seeded plates preincubated at appropriate temperature for 2 hours.

Three TIR1 expressing transgenes were tested for efficiency of germine specific removal of LAG-1; TIR1 expressed from the *sun-1* promoter [37], the *pie-1* promoter [71], or the *gld-1* promoter [37]. Efficacy of LAG-1 degradation was assessed by determining the relative mRNA abundance of GLP-1/LAG-1 transcriptional targets, *lst-1* & *sygl-1*, by qRT-PCR, following auxin treatment. We found that *gld-1* promoter driven TIR1 transgene (*ieSi64[gld-1p::tir1]*) gave the largest reduction of *lst-1* and *sygl-1* mRNA, and was used in this study.

### Immunostaining and Progenitor Zone size

The gonad dissection and staining procedure is as described [72]. The primary antibodies used are: WAPL-1 at 1:2000 (Rabbit, Novus, #49300002) [73], HIM-3 at 1:600 (Rabbit, Novus, 53470002), CYE-1 at 1:100 (mouse) [74], and HA at 1:100 (Rat, Roche, #11867423001), V5 at 1:1000 (mouse, Bio-rad, #MCA1360) [23], OLLAS at 1:500 (Rat, Novus, #NBP1-96713) [23]. Both CYE-1 and WAPL-1 antibodies staining gives similar estimate of progenitor zone size compared to REC-8 in *C. elegans* germline [73]. Hyperstack images were captured and a slice in mid-surface nuclei usually gives the best signal for both antibodies. Therefore this slice was used to count the number of rows of cells that are positive for either antibody, as described previously [73].

### Protein quantification

In this work, a method was developed to quantify protein level to study LAG-1, FBF-2, LST-1 and SYGL-1 accumulation changes in various genetic backgrounds. This method follows the approach of Brenner & Schedl (2016) [72] with following differences.

Quantification of LAG-1 level was performed with mid-L4 stage dissected gonads as LAG-1 accumulation was less variable than in 1-day old adults and this is the stage where LAG-1 ChIP-seq was performed. FBF-2 quantification is carried out in young-adult germlines (∼8 hrs post-L4/adult molt). At this stage, the wild type worms have started laying eggs and in tumorous animals (e.g., *gld-1* null germlines), the proximal tumor is still small enough to have not invaded the entire gonad; thus, the polarity of the germline is still intact [73]. Auxin treatment of animals carrying tagged LST-1 and SYGL-1 was started at mid-L4 stage and dissection and staining, thus, quantification of protein was carried out at 2 or 4 hours later at 25°C.

Apart from the protein to be quantified, gonads were co-stained with DAPI and antibody against WAPL-1. DAPI stained nuclei were used to count and mark cell diameter (cd) and WAPL-1 was used to measure progenitor zone length. WAPL-1 staining was also used to distinguish between different genotypes that were dissected and stained together. Hyperstack images were captured using a 63X objective lens on a spinning disk confocal microscope (PerkinElmer-Cetus, Norwalk, CT). Exposure time, which is kept constant for an individual experiment, is set by using auto exposure in Volocity software (Perkin-Elmer) for each experiment using an epitope-tagged strain in wild type background. To capture distal the end of germline, two overlapping hyperstack images were obtained to give a coverage length of ∼120-150 microns (equivalent to ∼40-50 cd). The images were exported, stitched in pairs, using Bio-Formats and Stitching plugin in Fiji [75–77]. In Fiji, DAPI images were used to draw a line, starting at the distal end to the desired cd [72].

Quantification of LAG-1, SYGL-1 and LST-1 was done for the distal most 25 cd, while for FBF-2 it was the distal most 35 cd. A line was drawn, in Fiji, using the DAPI image, starting at the distal tip and extending to the desired cd distance. A width of 75 pixels was used to collect fluorescent intensity for each line. Some of the gonads become out of focus across their length, therefore intensity values were collected for two or more slices of the hyperstacks for every pixel and a maximum value is selected for every pixel of the line. After manually drawing the line, ImageJ API (https://imagej.nih.gov/ij/developer/api/index.html) was used with custom python scripts to collect and store intensity data. The pixel intensities thus obtained, were processed similar to Brenner & Schedl (2016) to produce protein level graphs. Brenner & Schedl, *2016,* quantified GLD-1 levels using antibody against GLD-1, and used GLD-1 staining in *gld-1* null germlines to remove “non-specific signal/background noise”. Since protein level quantification in this study involved antibody staining against epitope-tagged proteins we instead used staining in N2, lacking the epitope tagged protein, to remove non-specific signal. Additionally, instead of using *spo-11(-)* germlines as internal normalization controls, we used strains carrying epitope-tagged protein, in an otherwise wild type background, as the internal control and have termed them as wild type germlines.

### mRNA detection and quantification

The procedure for *in situ* hybridization was adapted from Jones et al. 1996 [78]. Briefly, young adult animals were dissected in a glass dish. The dissected gonads were fixed with 3% paraformaldehyde / 0.25% glutaraldehyde / 0.1 M K2HPO4 (pH7.2) for 2 hours at room temperature, followed by post-fixed with 100% methanol at −20°C. After three washes in phosphate buffered saline with 0.1% Tween (PBST hereafter) to remove residual methanol, the gonads were incubated with 50 µg/ml protease K in PBST for 30 mins, followed by 15 mins re-fixation in 3% paraformaldehyde / 0.25% glutaraldehyde / 0.1 M K2HPO4 (pH7.2), three washes in PBST, 15 mins incubation in PBST with 2 mg/ml glycine to remove residue aldehyde, then another three washes in PBST prior to hybridization. The hybridization buffer contains: 5xSSC, 50% deionized formamide, 100 µg/ml autoclaved Herring sperm DNA, 50 µg/ml Heparin and 0.1 % Tween-20. The gonads were pre-hybridized in PBST/hybridization buffer (1:1) for 5 mins at 48°C, followed by 1 hour of incubation in hybridization buffer. PCR primers (see **S2 Table** for oligonucleotide information) were used to amplify *sygl-1* and *lst-1* cDNA from total RNA preparation and 100 ng of DNA was used to perform 35 rounds of single oligo PCR to incorporate digoxigenin labelled nucleotide into single stranded DNA probes that’s complementary to *sygl-1* and *lst-1* mRNA. Each probe was diluted in 1 ml hybridization buffer (for ten 100 µl hybridization). After O/N incubation in either probe, gonads were washed with hybridization buffer three times to remove the excess probe, followed by three washes in PBST. The gonads were then incubated with anti-digoxigenin antibody (Sigma, #11333089001) O/N at 4°C, followed by three washes in PBST, then developed in AP substrate BCIP/NBT (Sigma, #B5655). The detection of *sygl-1* takes 30mins and *lst-1* takes 1 hour.

Single molecule fluorescent *in situ* hybridization (smFISH) experiments were performed as described [43]. *lag-1* Stellaris smFISH probes were designed by and obtained from Biosearch Technologies. The fixed dissected gonads were incubated with probe at final concentration of 5 µM, at 37°C O/N. All the subsequent washes were done at 37°C: three washes in 2xSSC with 10% formamide, three washes in PBST. DAPI was introduced in the last wash to stain DNA. Gonads were mounted and imaged similar to immunostaining.

To quantify smFISH foci, hyperstack images of the distal end of the gonads were acquired using a 63X objective lens on a spinning disk confocal microscope (PerkinElmer-Cetus, Norwalk, CT). A suitable z-stack distance (0.4 micron) was used in order to capture all smFISH foci. Image acquisition along the entire thickness of the gonad results in progressive bleaching of the smFISH foci, hampering its accurate quantification. To circumvent this issue, instead of covering the entire thickness of the germline, we only took images of the germline covering one germ cell thickness from the surface. Two overlapping hyperstack images were acquired for each gonadal arm to get at least 25 cell diameters from the distal end. Images were stitched and further processed in Fiji. For every gonadal image, all the other external artifacts were cleared and gonads were rotated and cropped to get an optimum size to minimize the computational cost of further processing. 3D Objects Counter plugin [79] was used to get smFISH foci and their positions. A threshold value was selected, which would include most of the foci. For every germline, the first 25 cell diameters were marked, forming an imaginary line passing through the center of the gonadal tube. These markers and imaginary lines with a line-width of 100 pixels, roughly corresponding to about 10 microns, were used to assign smFISH foci to a particular cell diameter.

### Chromatin Immunoprecipitation (ChIP)

The overall ChIP procedure was adapted from Berkseth et al (2013) [80]. L4 stages animals raised at 20°C were used for both whole worm and germline specific LAG-1 ChIP experiments. Typically, freshly starved plates with abundant L1 stage animals were chunked onto NA22 bacteria plates [80], and grown to adult stage at 20°C. Gravid adult hermaphrodites were bleached to obtain a synchronized L1 population [80], which were plated onto NA22 plates (NA22 bacteria plates were seeded three days prior). Each NA22 plate (100 mm x 15 mm) housed around 5×10^4^ animals till L4 stages or 2×10^4^ animals till adult stage. If required, repeated growing/bleaching was used to obtain even larger quantities of animals. The L4 animals were washed off the plate using PBST and were further washed three times with PBST to remove NA22 bacteria, and frozen into small worm balls in liquid nitrogen.

The frozen worm balls were ground up in liquid nitrogen with a pre-cooled mortar and pestle. The worm powder was fixed in 1% paraformaldehyde in PBS for 15 mins, and then post-fixed with 125 mM glycine for 5 mins. After centrifugation, the worm pellet was washed in PBS three times before re-suspending in 1% SDS buffer. The worm suspension containing fixed DNA-protein complexes was sheared using a Bioruptor (Diagenode, Denville, NJ) on the high setting for 20 cycles (30 seconds ON/30 seconds OFF), to generate ∼ 500 base pair length fragmented DNA. After centrifugation, the supernatant was pre-cleared with protein-G Dynabeads (ThermoFisher, Waltham, MA) before the immunoprecipitation experiment.

For immunoprecipitation of LAG-1 from the whole worm L4 lysate, different GFP and FLAG antibodies were first tested for their ability to immunoprecipitate LAG-1 (**S4B Fig**). Two antibodies were chosen for further analysis: FLAG M2, which has have been successfully used in other ChIP experiments [81, 82], and anti-GFP antibody, which immunoprecipitated the highest amount of LAG-1 (**S4B Fig**). Two µg of either FLAG (mouse, Sigma, #F1804) or GFP (goat, Rockland, #600-101-215) antibodies was used to immunoprecipitate LAG-1 protein from 2 grams of worm powder. After O/N incubation at 4°C, the precipitated DNA-protein complexes were washed and eluted in 1% SDS for 10 mins. DNA was then reverse crosslinked to prepare sequencing libraries following the manufacturer’s instructions (Kapa Biosystem, #KK8500). To immunoprecipitate germline LAG-1, L4 animals were used as this stage lacks LAG-1 from late stages of oogenesis that are not functioning in GLP-1 dependent stem cell fate regulation (**Fig 2**). However, direct ChIP with streptavidin beads was not successful, likely due to interference from large amounts of endogenous biotinylated protein [54, 55]; hence, a sequential ChIP method was developed to overcome this issue. First, 300 µl FLAG M2 beads/6 grams of worm powder per replicate (Sigma, #M8823) was used to precipitate all LAG-1 from both germline and somatic tissues, following the same procedure from whole worm ChIP. After eluting in 1% SDS, the DNA-protein complexes was diluted five-fold and re-immunoprecipitated with streptavidin beads to specifically pull-down biotinylated germline LAG-1. Since the binding between biotin and streptavidin is very stable, the precipitated DNA-protein complex was washed in 2% SDS, then in 1% SDS, for 10 mins each, to remove any none specific binding. Unlike the whole worm ChIP, the DNA amount after sequential ChIP was ultra low, so the NGS libraries were prepared directly with DNA on the streptavidin beads following manufacturer’s instructions (Kapa Biosystem, #KK8500).

Sequencing and Bioinformatic analysis was performed by Genome Technology Access Center at the McDonnell Genome Institute (GTAC @ MGI, Washington University in St. Louis). Short sequencing reads (50 nt) were aligned back to *C. elegans* reference genome and ChIP-seq peaks were identified using MACS2.0 [83] by comparing ChIP library to input DNA library, for both whole worm and germline ChIP experiments. The association of peak to closest gene annotation was done with HOMER suite [46]. HOMER suite was also used to identify overly represented motifs in ChIP-seq data. Gene list comparisons and Venn diagrams preparation were both conducted in R.

### RNA-seq analysis

All RNA-seq experiments performed in this study used total RNA from dissected gonads. All strains carried *gld-2 gld-1* double mutant background. For GLP-1 ON and OFF RNA-seq, gain of function (gf) allele *glp-1(ar202)* or null allele *glp-1(q175)* were introduced into the *gld-2(q497) gld-1(q485)* mutant background. *glp-1(ar202)* is a temperature sensitive gain of function allele, used to produce cells throughout the germline that are undergoing GLP-1 signaling. This increased our ability to detect gene expression differences. For the auxin induced LAG-1 degradation time course RNA-seq experiments, *lag-1(oz536oz537)* and *ieSi64[gld-1p::tir1]* were introduced into *gld-2(q497) gld-1(q485); glp-1(ar202)*. Animals were raised at 25°C for 48 hours from synchronized L1s population before dissection. For shorter periods of auxin treatment (e.g, 2 hours and 4 hours), animals grown on NGM plates were transferred to seeded auxin plates (pre-conditioned to 25°C) and treated for the required time, prior to gonad extrusion.

Fifty young adults were dissected in a glass dish and the gonads were dissected away from rest of the animal’s body at the spermatheca. The isolated gonads were centrifuged for 15 sec and re-suspended in 500 µl Trizol (Invitrogen, #15596026). The RNA isolation was done according to manufacturer’s instructions. RNA was then treated with DNase (Thermo Fisher Scientific, #EN0525) to remove the contaminating genomic DNA before proceeding to the removal of ribosome RNA (rRNA). The rRNAs were removed with the Ribo-Zero kit (Illumina, #RZE1206) according to manufacturer’s instructions. The conversion of RNA to dsDNA was done with NEBNext RNA first strand (NEB, #E7525S) and second strand (NEB, E6111S) kits. The converted dsDNA was used to prepare NGS libraries similar to ChIP-seq (Kapa Biosystem, #KK8500).

The sequencing and bioinformatic analysis was performed by the Genome Technology Access Center at the McDonnell Genome Institute. The samples were sequenced on an Illumina HiSeq 3000 with 50 nt reads to a mean depth of 34.5 million reads and then aligned back to the Ensembl Release 76 *C. elegans* genome with STAR version 2.0.4b [84]. Gene counts were then quantitated with Subread:featureCount version 1.4.5 [85]. The gene counts were then TMM scaled with EdgeR [86] and voomWithQualityWeights [87] transformed with Limma [88]. A Limma generalized linear model without intercept was then fitted to the data and tested for statistical significance via Limma’s empirical Bayes moderated t-statistics. The genes were considered to be activated by GLP-1 if the following three criteria were met: (1) its expression in GLP-1 ON gonads was at least 2.0 fold compared to GLP-1 OFF gonads. (2) the Benjamini-Hochberg false discovery rate (FDR) is less than 0.05 and (3) the gene expression level as determined by Counts Per Million (CPM) must be greater than 2.0. The same criteria were applied to obtain the gene list for either activated or repressed by LAG-1 in the time course RNA-seq analysis. Gene list comparisons and Venn diagrams preparation were both conducted in R.

### Immunoblot analysis

The immunobloting was performed as described [78]. Immunoblot was used to determine the efficiency of different commercial antibodies in pull-down of LAG-1 in ChIP conditions (**S4 Fig**). After ChIP experiments, the protein DNA complexes were eluted in 1x SDS sample buffer by heating at 95°C for 5 mins. The eluted samples were run on 10 % SDS-PAGE gel, then transferred to PVDF membrane. To detect LAG-1 after ChIP, FLAG antibody (Sigma, F1804) was used at 1:1250 dilution. The detection of signal was achieved with HRP-conjugated anti-mouse secondary antibody (Jackson Immunoresearch, #115-035-146).

### qRT-PCR analysis

For analysis of mRNA abundance in tumorous germline of relevant genotypes (**Fig 5, 7**), total mRNA was isolated from dissected gonads with Trizol (Invitrogen, #15596026) following manufacturer’s protocol. The RNA was then DNase treated before cDNA synthesis with Superscript III (Invitrogen, #18080051). The cDNAs were usually diluted five fold prior to qPCR analysis. For analysis of DNA enrichment after ChIP experiments, the DNA concentration was determined and the same amount of DNA was used for each qPCR reaction. iTaq Universal SYBR Green Supermix (Biorad, #1725121) and CFX96 Real time System (Biorad) were used for qPCR analysis. The delta Ct method was used to calculate the relative mRNA abundance compare to the *ama-1* control [89]. The primers used for qPCR analysis are listed in **S2 Table**.

### Data processing and statistical analysis

Data stored as either text or csv files, were imported into an R programming environment, and graphs were plotted using the ggplot2 package (http://ggplot2.org). The significance bars were generated using the ggsignif package (https://CRAN.R-project.org/package=ggsignif) in R. Statistical significance was determined using either two-tailed Student’s t-test (for n < 30) or Z-test (for n ≥ 30).

## Acknowledgments

We thank our colleagues in worm community for unpublished alleles: Iva Greenwald, Columbia University, *lag-1(ar611);* William Kelly, Emory University, *ckSi11*. We also thank Julie Ni and Sam Gu for their help in the initial phases of ChIP-seq analysis and our colleagues Mike Nonet, Jane Hubbard, Iva Greenwald, Dave Hansen, David Greenstein, Jordan Ward, and Dan Dickenson for helpful advice and discussions. We thank WormBase, funded by the National Human Genome Research Institute; the Caenorhabditis Genetics Center, funded by National Institutes of Health Office of Research Infrastructure Programs; and Shohei Mitani and the Japanese National Bioresource Project for strains. This work was supported by National Institutes of Health R01 GM-100756 to T.S.

## Figure legends

**Supplemental figure 1.**
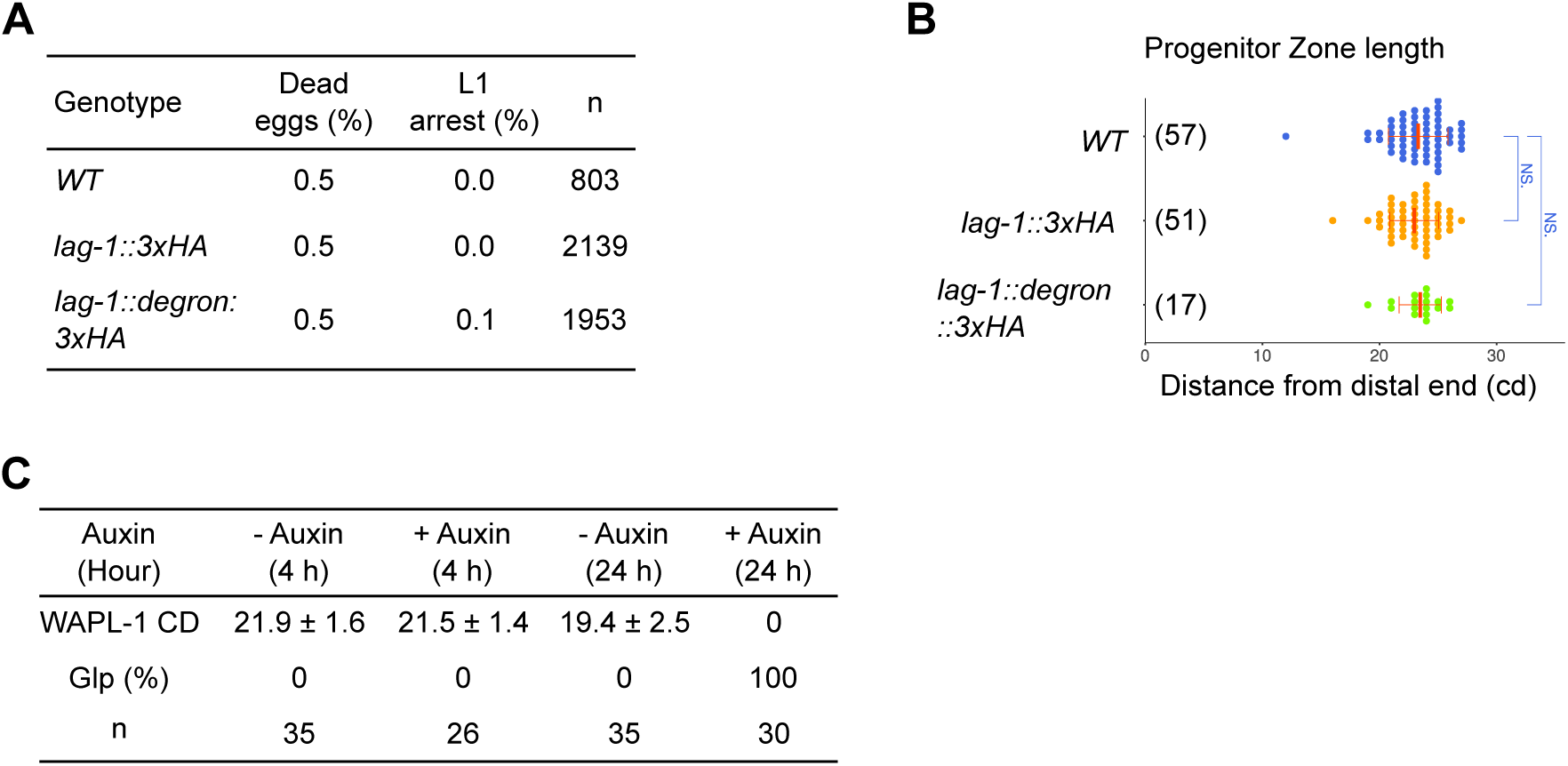
Characterization of epitope-tagged *lag-1* alleles. (A & B) The dead eggs and L1 arrest frequency (A) and progenitor zone length (B) in WT, *lag-1(oz530[lag-1::3xHA])* and *lag-1(oz536oz537[lag-1::degron::3xHA])*. (B) Graph showing progenitor zone length, in cell diameters, between the distal tip of the germline and the row of cells at proximal end of the continuous zone of WAPL-1 staining for L4 hermaphrodites of indicated genotype. Data are plotted as horizontal dot plots with each dot representing length in cell diameter to zone end for one gonad. Numbers in bracket shows the sample size. Thick vertical lines represent mean and horizontal lines represent mean ± SD. P-value ≤ 0.01 (*); ≤ 0.001 (**); ≤ 0.0001 (***); > 0.01 non-significant (NS.). (C) Progenitor zone length and premature meiotic entry of progenitor zone cells (Glp) phenotype after L4 stage animals were treated with or without auxin for either 4 hrs or 24 hrs (in Fig 2D & E). The genotype for auxin treatment was *lag-1(oz536oz537[lag-1::degron::3xHA]); ieSi64[gld-1p::TIR1::mRuby::gld-1 3’UTR]*. The *lag-1::3xHA* and *lag-1::degron::3xHA* strains appears phenotypically wild type; we did not observe phenotypes associated with *lag-1* loss of function [11, 18], including loss of *glp-1* - embryonic lethality or a smaller progenitor zone due to stem cells undergoing spatially premature meiotic entry, loss of *lin-12* - egg-laying/vulva defects, or loss of both *lin-12* and *glp-1* - Lag larval arrest.

**Supplemental figure 2.**
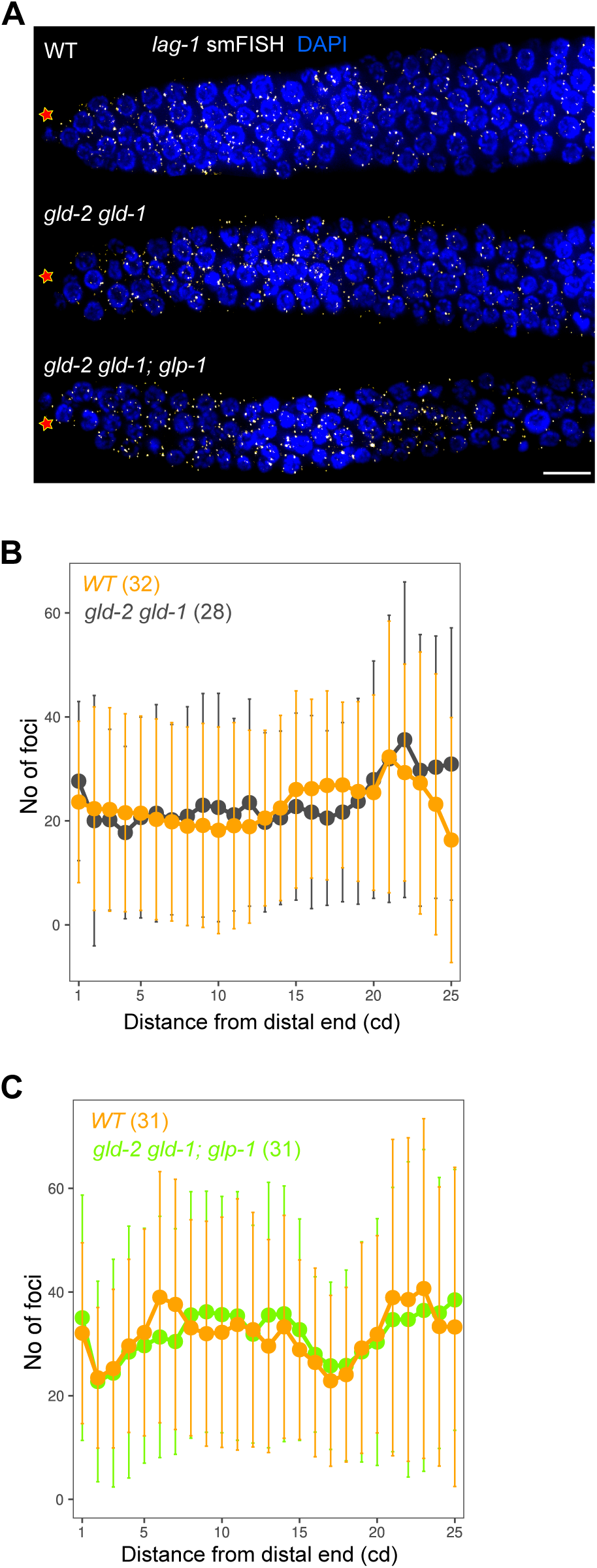
*lag-1* mRNA expression analysis by smFISH. (A) Z project of one nucleus thick (see **Materials and Methods**) distal germlines of L4 hermaphrodites, probed for LAG-1 transcripts using smFISH (white) and DAPI (blue). Asterisk, distal end; Scale bar, 10 μm. (B & C) Density plot of *lag-1* mRNA foci for indicated genotype. Numbers in bracket shows the sample size. Additionally, there was no difference in *lag-1* mRNA level from GLP-1 ON versus GLP-1 OFF transcriptomics analysis.

**Supplemental figure 3.**
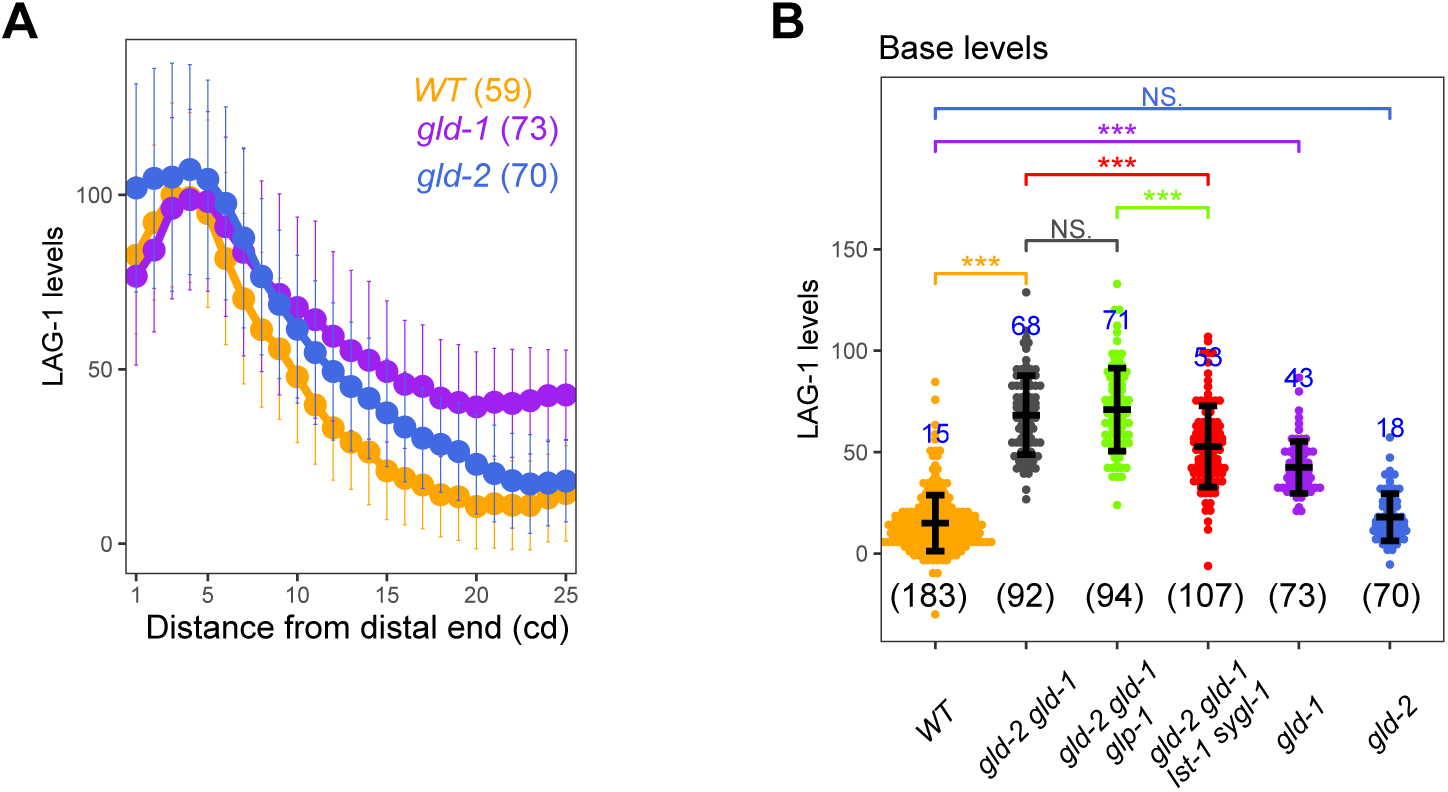
LAG-1 levels analysis. (A - B) Plot of LAG-1 levels (A) and comparison of LAG-1 base (B) for indicated genotype. *lag-1(oz530[lag-1::3xHA])* was used for quantification; See **S1 Table** for the complete genotypes. Numbers indicate mean values of LAG-1 level for each genotype and numbers in bracket shows the sample size. Dots, mean (A) or data points (B); Error bars, mean ± SD. P-value ≤ 0.01 (*); ≤ 0.001 (**); ≤ 0.0001 (***); > 0.01 non-significant (NS.).

**Supplemental figure 4.**
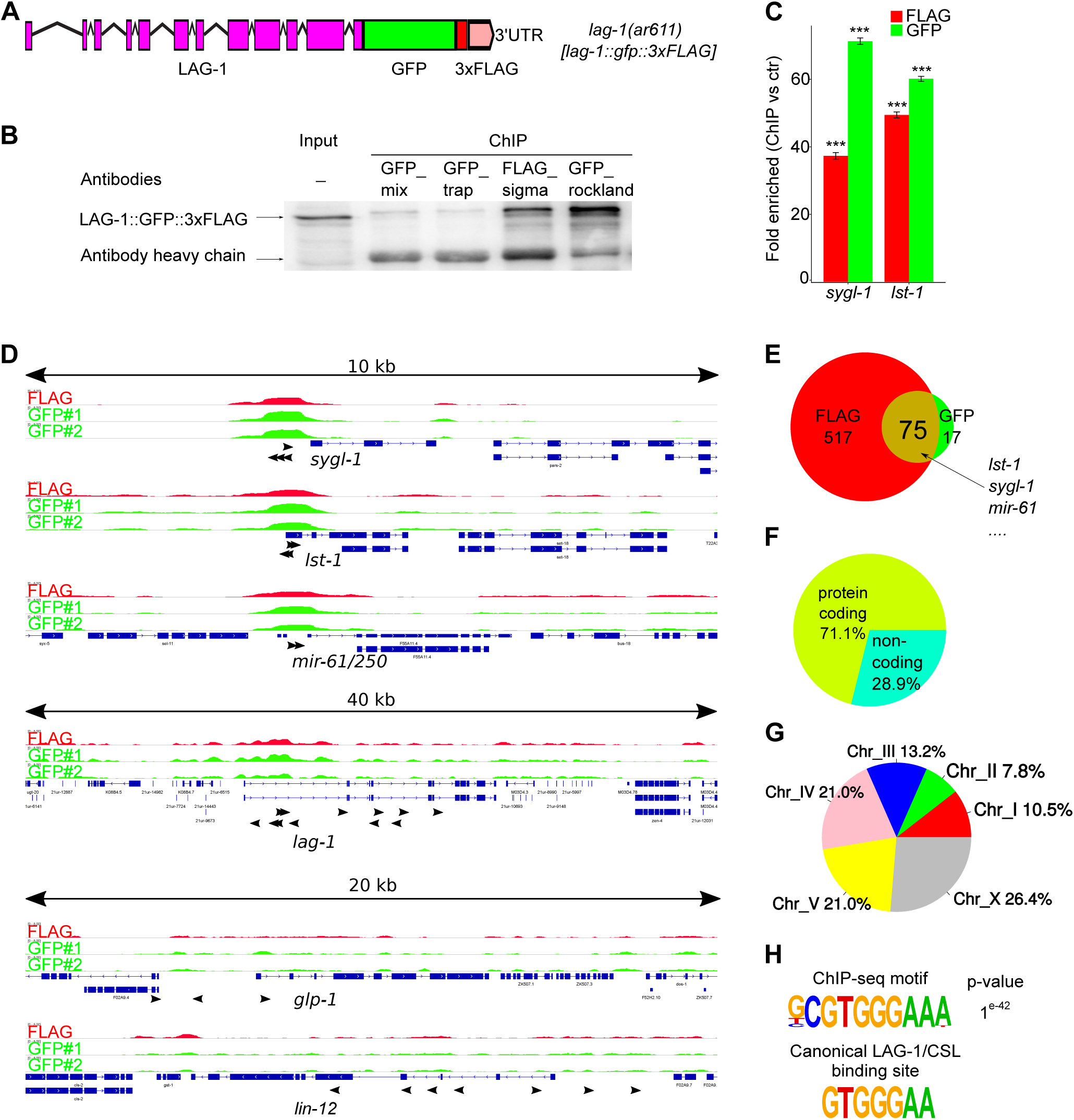
Genome-wide identification for LAG-1 targets in whole animal by ChIP-seq analysis. (A) Diagram of *lag-1* allele at endogenous locus, *lag-1(ar611[lag-1::gfp::3xFLAG])* (gift from Iva Greenwald), used for whole worm ChIP-seq analysis. (B) Different antibodies were tested to determine if they are competent for ChIP. Same amount of extracts were used for each ChIP experiment, followed by western blot analysis with FLAG antibody. Bottom band detects the heavy chain of antibodies. FLAG antibody from Sigma and GFP antibody from Rockland were selected for the analysis below. (C) ChIP-qPCR analysis for *sygl-1* and *lst-1* promoter regions bound by LAG-1. A non-peak region in the *xol-1* promoter was used as a negative control (ctr) and set as 1. *** for p<0.0001. Error bars, mean ± SD. (D) Genome browser tracks showing 10 kb genomic region for *sygl-1*, *lst-1 and mir-61/250,* 40 kb genomic region for *lag-1*, and 20 kb genomic region for *glp-1 and lin-12* after ChIP-seq. Raw reads were normalized to control, and signal intensity were presented as log2 fold change. Black arrow heads, canonical LAG-1/CSL binding motif GTGGGAA [17,19,20]. (E) Venn diagram showing the overlapping genes identified through FLAG antibody and GFP antibody ChIP-seq analysis. Both data lists were filtered for more than 2-fold change of signal (ChIP/control) with a moderate False Discovery Rate (FDR<0.05). (F & G) Protein coding vs. non-coding distribution (F) and chromosome position (G) of 75 genes from E. The overly represented motif discovered by HOMER suite with the ChIP-seq data (top) and the canonical LAG-1/CSL binding sequence [17,19,20](bottom). See **S3 Table**.

**Supplemental figure 5.**
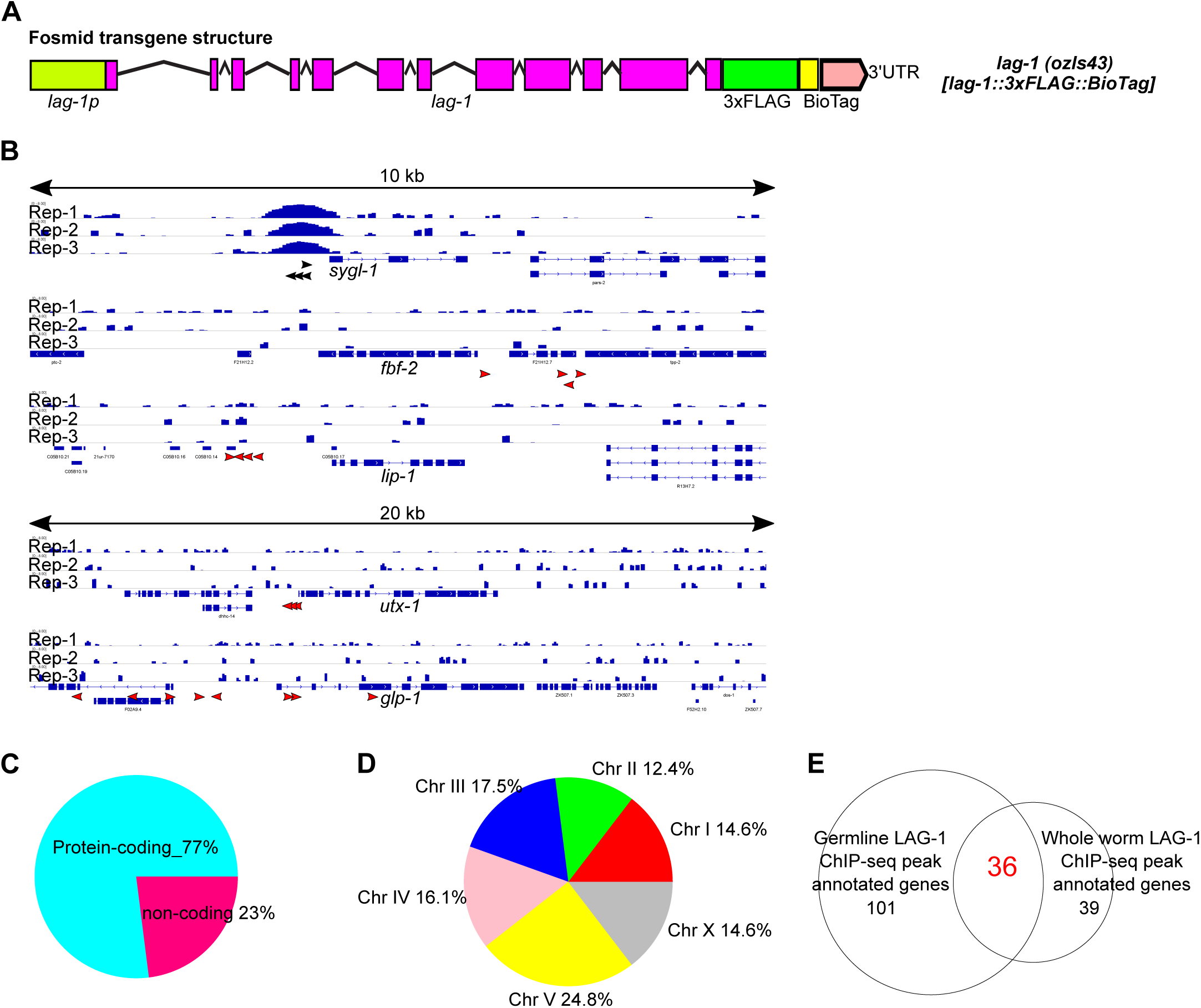
Supplemental information for genome-wide identification of germline LAG-1 targets. (A) Diagram of *lag-1(ozIs43[lag-1p::lag-1::3xFLAG::BioTag::lag-1 3’UTR])* fosmid transgene. 3xFLAG and BioTag sequence were inserted into a fosmid that contains native regulatory sequence for *lag-1* gene. *lag-1(ozIs43)* was able to rescue *lag-1* deletion allele *tm3052*. (B) Genome browser tracks showing *sygl-1* and other four putative germline GLP-1/LAG-1 transcriptional targets from literature: 10 kb genomic region for *sygl-1* (this study, [21]), *fbf-2* [30] *and lip-1* [31]*, 2*0 kb genomic region for *utx-1* [32] and *glp-1* [17]. Raw reads were normalized to control, and the signal intensity were presented as log2 fold change. Black arrow heads, canonical LAG-1/CSL binding motif GTGGGAA [17,19,20]. Red arrow heads, LAG-1 binding site (LBS) from original references where LAG-1 was suggested to bind. (C & D) Protein coding/ non-coding distribution (C) and chromosome position (D) of 137 genes from Fig 4D. (E) Venn diagram showing the overlapping genes from germline and whole worm ChIP-seq analysis of LAG-1. See **S4 Table**.

**Supplemental figure 6.**
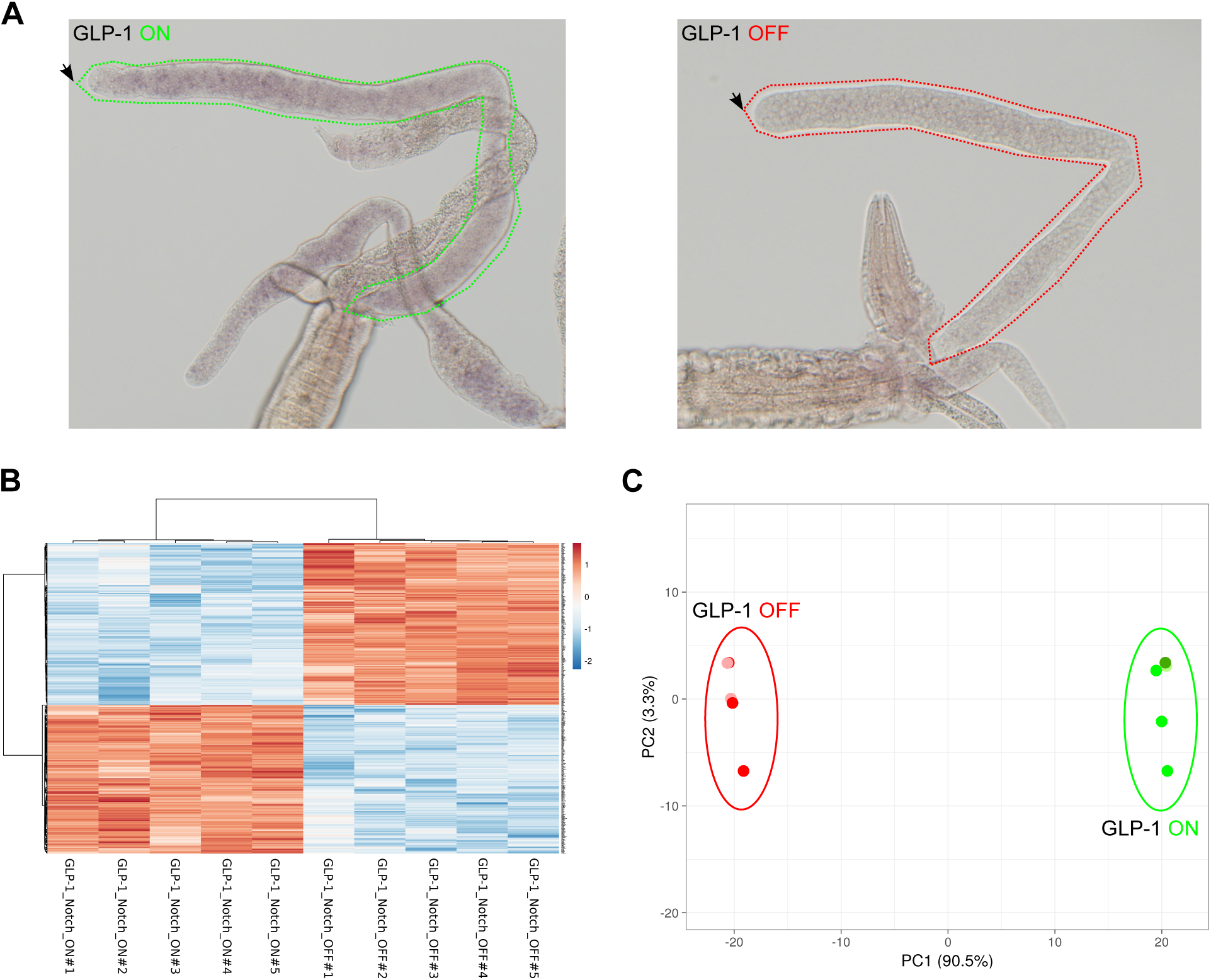
Supplemental information for transcriptomic analysis to identify GLP-1-dependent genes. (A) *In situ* hybridization used to determine *lst-1* mRNA expression in young adult animals. The genotypes are, **GLP-1 ON**: *gld-2(q497) gld-1(q485); glp-1(ar202)* and **GLP-1 OFF**: *gld-2(q497) gld-1(q485); glp-1(q175)*. Dotted lines outline the boundary of the gonad. Black arrow indicates the distal tip of the gonad. (B & C) Heatmap (B) and principle component analysis (PCA) (C) were generated using the top 500 genes with the most significant p-value from differential gene expression analysis.

**Supplemental figure 7.**
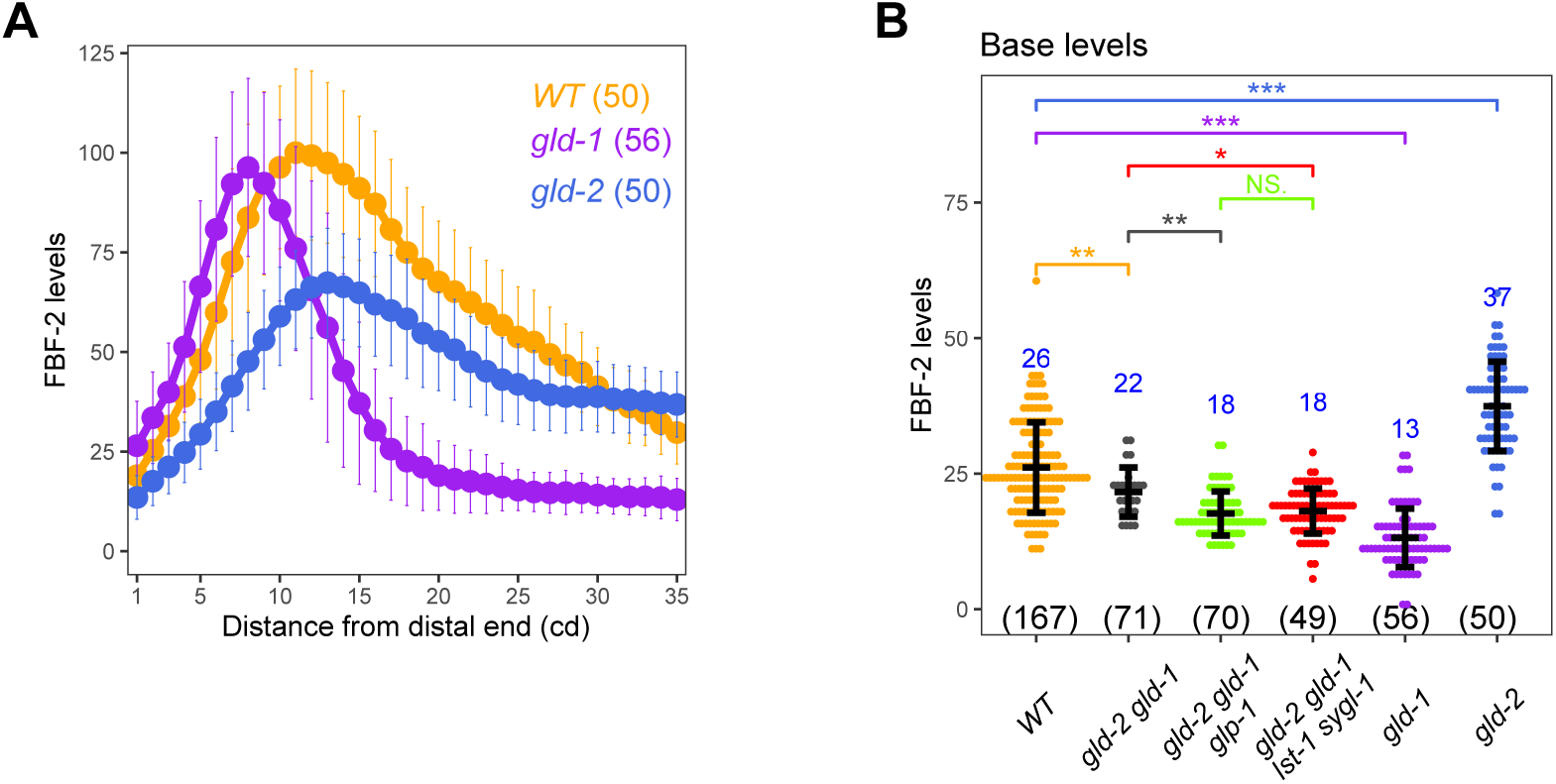
FBF-2 accumulation analysis. (A - B) Plot of FBF-2 levels (A) and comparison for FBF-2 base (B) for indicated genotype. *fbf-2(q932[3xV5::fbf-2])* is used for quantification. See **S1 Table** for the complete genotypes. Numbers indicate mean values of FBF-2 level for each genotype and numbers in bracket shows the sample size. Dots, mean (A) or data points (B); Error bars, mean ± SD. P-value ≤ 0.01 (*); ≤ 0.001 (**); ≤ 0.0001 (***); > 0.01 non-significant (NS.).

**Supplemental figure 8.**
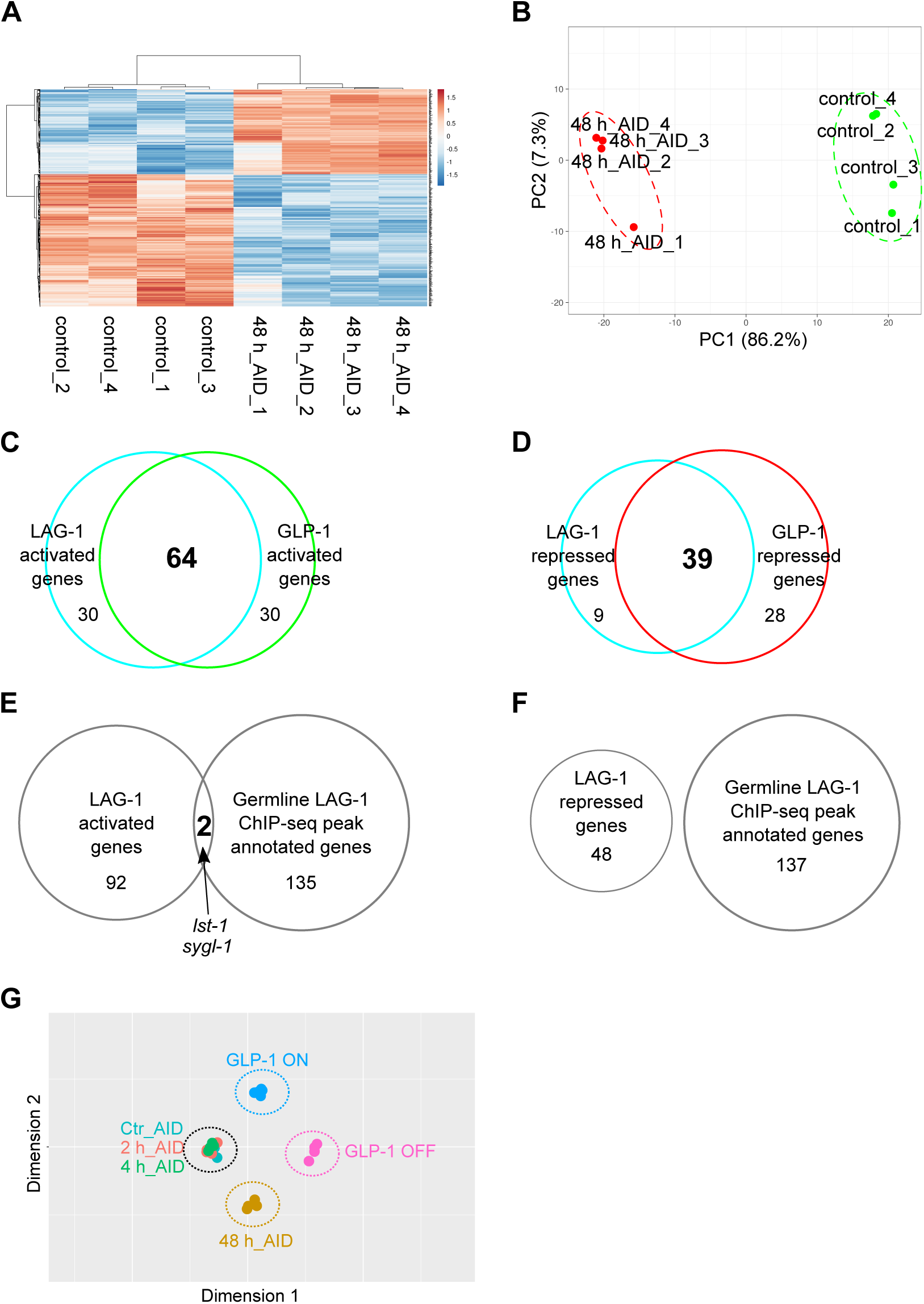
Time-course transcriptomic analysis upon LAG-1 degradation in germline. (A & B) Heatmap (A) and principle component analysis (PCA) (B) for top 500 genes with most significant p-values, with the differential gene expression analysis done between animals treated with or without auxin for 48 hours. (C & D) The differentially-expressed genes upon auxin treatment for 48 hours were compared to the differentially-expressed genes in GLP-1 ON vs. OFF to identify the overlapping genes activated (C) or repressed (D) by both LAG-1 and GLP-1. (E & F) The differentially-expressed genes upon auxin treatment for 48 hours were compared to putative LAG-1 targets through LAG-1 germline ChIP-seq analysis to determine the LAG-1 transcriptional targets (E) and if LAG-1 can repress gene expression (F). (G) Multiple dimensional scaling analyses showing the similarities of the RNA-seq samples conducted in this study. Five biological replicates each were conducted for GLP-1 ON (in blues circle) and GLP-1 OFF (in pink circle). Four biological replicates were conducted for time course RNA-seq analysis following LAG-1 degradation by auxin treatment. The 48-hour auxin treated samples were grouped in yellow circle. For the shorter period of treatment, 2 hours and 4 hours, the transcriptomic profile are similar to the untreated samples and were grouped in the black circle. See **S6-7 Tables**.

**Supplemental figure 9.**
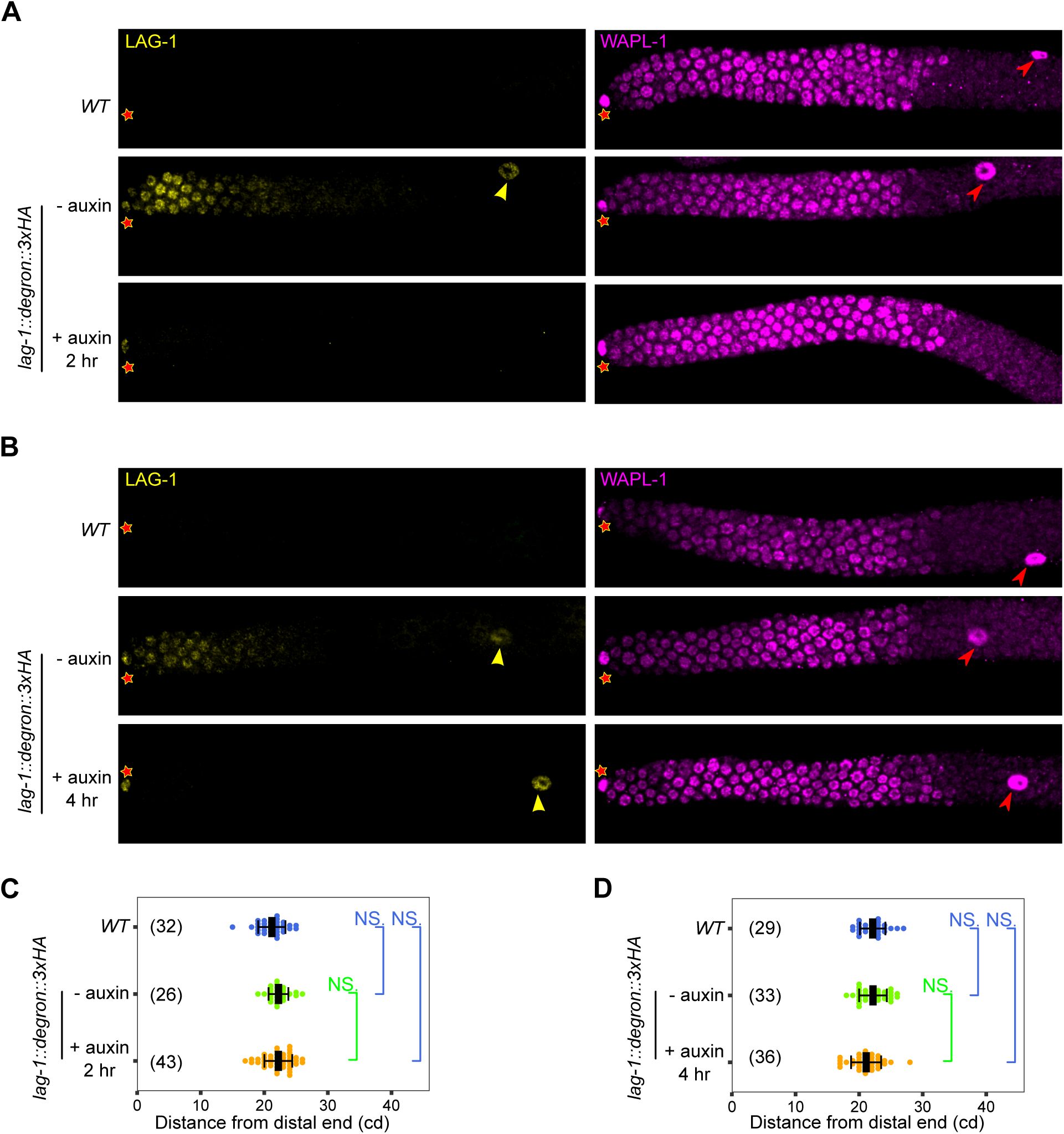
Time-course auxin treatment to degrade LAG-1. (A & B) Representative images of anti-HA-stained (LAG-1, yellow, left panels) and WAPL-1-stained (pink, right panels) after L4 stage animals were treated with or without auxin for 2 hours (A) or 4 hours (B) at 25°C. Asterisks, distal end of germline. Yellow arrowheads, LAG-1 accumulation in the sheath cells, pink arrowheads, WAPL-1 accumulation in sheath cells. (C & D) Graph showing distance, in cell diameters, between the distal end of the germline and the row of cells at proximal end of the continuous zone of WAPL-1 staining for auxin-treated hermaphrodites. Data are plotted as horizontal dot plots with each dot representing length in cell diameter to zone end for one gonad. Numbers in bracket shows the sample size. Thick vertical lines represent mean and horizontal lines represent mean ± SD. P-value ≤ 0.01 (*); ≤ 0.001 (**); ≤ 0.0001 (***); > 0.01 non-significant (NS.).

**Supplemental figure 10.**
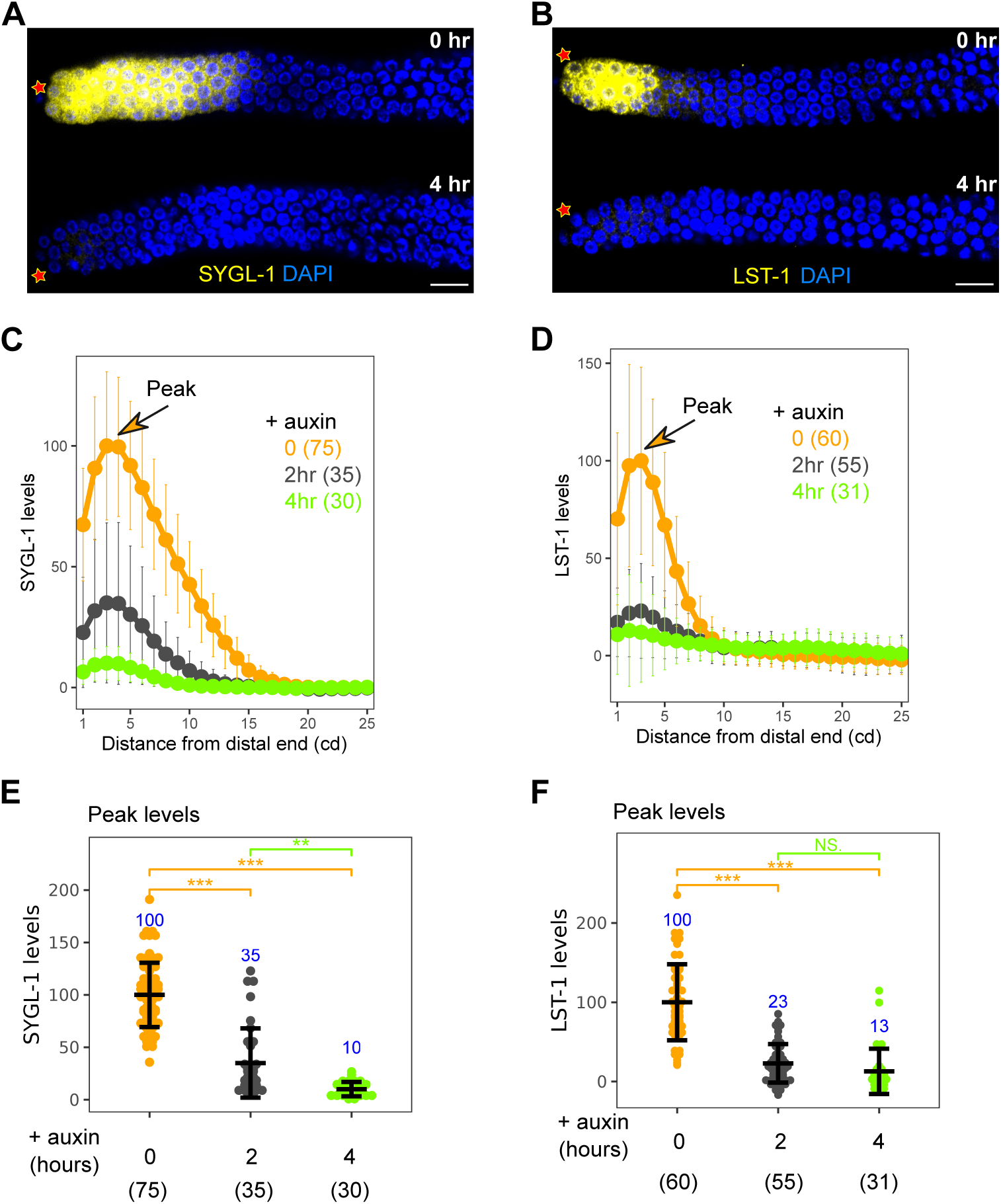
SYGL-1 and LST-1 accumulation analysis following 2-hour and 4-hour auxin treatment to degrade LAG-1 protein. (A & B) Representative images of SYGL-1 (A) and LST-1 (B) accumulation after animals were treated with or without auxin for 4 hours. The genotype for auxin treatment are *sygl-1(q983[3xOLLAS::sygl-1]); lag-1(oz536oz537[lag-1::degron::3xHA]); ieSi64[gld-1p::TIR1::mRuby::gld-1 3’UTR]* (for quantification of SYGL-1 levels) and *lst-1(q1003[lst-1::3xOLLAS])*; *lag-1(oz536oz537[lag-1::degron::3xHA]); ieSi64[gld-1p::TIR1::mRuby::gld-1 3’UTR]* (for quantification of LST-1 levels). Note that the images, taken at identical exposure time, were processed identically to make residual protein visible in the auxin treated germlines, this resulted in slightly saturated signal in the untreated germlines. (C & D) Plots of SYGL-1 (C) and LST-1 (D) levels in auxin-treated germlines. Numbers in bracket shows the sample size. Dots, mean. Error bars, mean ± SD. (E & F) Comparison of SYGL-1 (E) and LST-1 (F) peak levels (see C & D) in auxin-treated germlines. Numbers indicate mean values for each genotype and numbers in bracket shows the sample size. Dots, data points. Error bars, mean ± SD. P-value ≤ 0.01 (*); ≤ 0.001 (**); ≤ 0.0001 (***); > 0.01 non-significant (NS.).

**S1 Table.** Strains used in this study.

**S2 Table.** Oligonucleotides used in this study.

**S3 Table.** List of putative LAG-1 peak regions identified through whole worm ChIP-seq analysis.

**S4 Table.** List of putative LAG-1 peak region identified through germline ChIP-seq analysis.

**S5 Table.** List of genes activated or repressed by GLP-1.

**S6 Table.** List of genes activated or repressed by LAG-1, after 48 hours of auxin treatment.

**S7 Table.** List of genes activated or repressed by LAG-1, after 2 and 4 hours of auxin treatment.

